# Choreographed morphogenetic events underlie early foregut development in the mouse embryo

**DOI:** 10.64898/2026.08.21.746211

**Authors:** Jenny Kretzschmar, Guillermo Serrano Nájera, Sonia Agüera-Gonzalez, Henry Westmacott, Casper van Bavel, Pranay Shah, Lara Krasinska, Tom Smith, Rob Jelier, Katie McDole

## Abstract

Between embryonic days 7.5 and 8.5, the mouse embryo undergoes a dramatic rearrangement of its entire anterior, a process known as ventral folding: the near-simultaneous morphogenesis of the cardiac crescent, cranial headfolds, and anterior foregut that together establish the antero-ventral body plan. While cardiac morphogenesis has been studied in detail, the cellular and mechanical basis of foregut involution remains largely undefined. Using combined light-sheet and spinning-disk live imaging spanning the full window of foregut formation, we show that involution proceeds through a stereotyped morphological programme that does not require actomyosin contractility for its initiation. Additionally, involution is preceded by a spatiotemporally restricted wave of apoptosis in embryonic visceral endoderm (emVE) cells, which extrude bidirectionally. A lineage- and stage-resolved bulk RNA-sequencing of emVE, definitive endoderm, and epiblast populations identifies a differential-adhesion and cell-cycle signature underlying this behaviour. Revisiting a visceral-endoderm-specific *Bmp2* knock-out mutant, we find that BMP2 controls involution indirectly, by directing notochord positioning and thereby the geometry of the surrounding heart and headfold mechanics. Together, these findings reframe ventral folding as a single coordinated geometric process rather than a set of independent organ forming events.

## Introduction

The transformation of the early post-implantation mouse embryo from an egg cylinder into a complex, structured system with specialised tissues and organs is one of the most dramatic events in early mammalian development. Between embryonic days 7.5 to 8.5 (E7.5–E8.5), the embryo has established the first three germ layers and initiates organogenesis^1–3^. The brain, heart, and gut begin to form within this brief window in close proximity on the anterior of the embryo^4–6^. The morphogenesis of each is closely coordinated in both timing and geometry. How this coordination is achieved at the level of tissue mechanics, cell behaviour, and molecular signalling remains largely unresolved.

The anterior side of the embryo is the first region to acquire definitive form and function. Within hours of their specification, nascent cardiomyocytes in the cardiac crescent coalesce and begin to contract, producing the first beats of the linear heart tube as it assembles^6,7^. Immediately dorsal to this nascent heart, the cranial neural plate expands and elevates to generate the headfolds, or early brain^4,8,9^. Just ventral to both on the surface of the embryo, the anterior endoderm undergoes a dramatic inward fold that generates the foregut pocket, the precursor of the foregut tube. These three near-simultaneous morphological events together constitute the process known as ventral folding^4,5,10^.

Despite the central role of ventral folding in early mammalian embryogenesis, our understanding of the cellular and molecular events surrounding this process is incomplete. Early heart development has been mapped with exquisite molecular and cellular resolution, with transcriptional profiles, lineage hierarchies, and live-imaging studies now describing the specification, migration, and assembly of the cardiac crescent in considerable detail^7,11,12^. The formation of the early gastrointestinal system at these stages, by contrast, has received far less attention. The foregut undergoes a dramatic shape change in the early embryo: a squamous sheet of endoderm on the anterior surface suddenly and rapidly involutes and extends inward, ultimately giving rise to the pharynx, thyroid, lungs, liver, gall bladder, pancreas, stomach, and upper gastrointestinal tract^13,14^. The most detailed prior anatomical account of this process comes from Madabhushi and Lacy, who showed that visceral-endoderm-specific deletion of *Bmp2* produces a disorganised anterior phenotype with mispositioned heart and headfolds and disrupts ventral folding^5,10^. Their work established the visceral endoderm (VE) as an essential source of signalling activity for anterior organogenesis, but how the foregut itself involutes, and how that involution is coordinated with the surrounding tissues remains unclear.

Complicating our understanding of foregut involution is the unusual cellular composition of the anterior gut endoderm. At E7.5 the surface endoderm is a mosaic of two developmentally distinct populations. Embryonic visceral endoderm (emVE) cells are descended from the primitive endoderm of the blastocyst and, although extraembryonic in origin, contribute substantially to the embryo proper, lining the egg cylinder before gastrulation and persisting within the gut endoderm long after tube formation^15–17^. Definitive endoderm (DE) cells, by contrast, are specified from the epiblast during gastrulation, ingress through the primitive streak, and intercalate widely into the overlying VE layer, dispersing the emVE into a salt-and-pepper pattern^16,18^. The two lineages converge into a single epithelial sheet but retain distinct molecular identities. Within the VE, an earlier specified subpopulation, the anterior visceral endoderm (AVE), patterns the antero-posterior axis through localised secretion of Nodal and Wnt antagonists^3,19,20^. The anterior endoderm at the site of foregut involution is therefore a tissue comprised of an established signalling history and a heterogeneous cellular composition, raising the possibility that emVE and DE cells contribute differently to the morphogenetic programme that follows.

The largest hurdle to generating a complete cellular and mechanistic picture of ventral folding has, in large part, been hindered by the difficulty in visualising the embryo live during these stages. The mouse embryo at E7.5 develops *in utero*, is sensitive to perturbation, and undergoes substantial growth over the few hours during which the foregut forms - features that have until recently rendered it intractable to long-term live imaging in conventional microscope systems. As a result, while fixed snapshots and end-stage phenotypes exist, very little is known about how the dynamic cellular behaviours within the anterior endoderm are coordinated with the elevation of the headfolds and the convergence of the cardiac crescent to produce the stereotyped geometry of the involuting foregut. The complex cellular choreography of foregut involution itself has never been described.

Here we combine adaptive light-sheet microscopy^21^, customised spinning-disk imaging, fluorescent reporter mouse lines, bulk RNA sequencing, chemical and genetic perturbations, and 3D morphometrics, to provide the first comprehensive tissue-, cellular-, and molecular-level characterisation of foregut involution in the mouse embryo. We define a stereotyped morphological progression of foregut formation across E7 to E8 day old embryos using light-sheet microscopy and FlowShape^22^, which provides a quantitative framework to track embryo shape over time. This allows us to reconstruct the morphogenesis of the embryo over the involution stages and align embryos across space and time. We identify a spatiotemporally restricted apoptotic cell death event that is exclusive to anterior emVE cells and tightly correlated with the onset of invagination. We show that emVE cells extrude bidirectionally through the assembly of a contractile actin cable. We complement these observations with a lineage-separated, stage-resolved bulk RNA-sequencing dataset that distinguishes anterior and posterior emVE, DE, and mesoderm/epiblast across the involution window. Finally, by reexamining the VE-specific *Bmp2* knockout, we uncover a previously unappreciated requirement for proper notochord positioning in elevating the neural headfolds, supporting a model in which ventral folding emerges from the coupled mechanics of foregut, headfold, and cardiac morphogenesis rather than from any one of these tissues alone.

## Results

### Foregut involution proceeds through a stereotypic morphological programme

To resolve the dynamics of foregut involution at cellular resolution, we performed long-term live imaging of E7.5 mouse embryos using both adaptive light-sheet microscopy^21^ and spinning-disk microscopy with custom-built imaging chambers. Embryos expressing various nuclear, membrane, VE- or DE-specific reporters were imaged from the early bud (EB, approximately E7.5) stage through to the late headfold (LHF, approximately E8.0) stage, capturing the entirety of foregut formation (Figure 1A, S1A; Movie S1).

**Figure 1:**
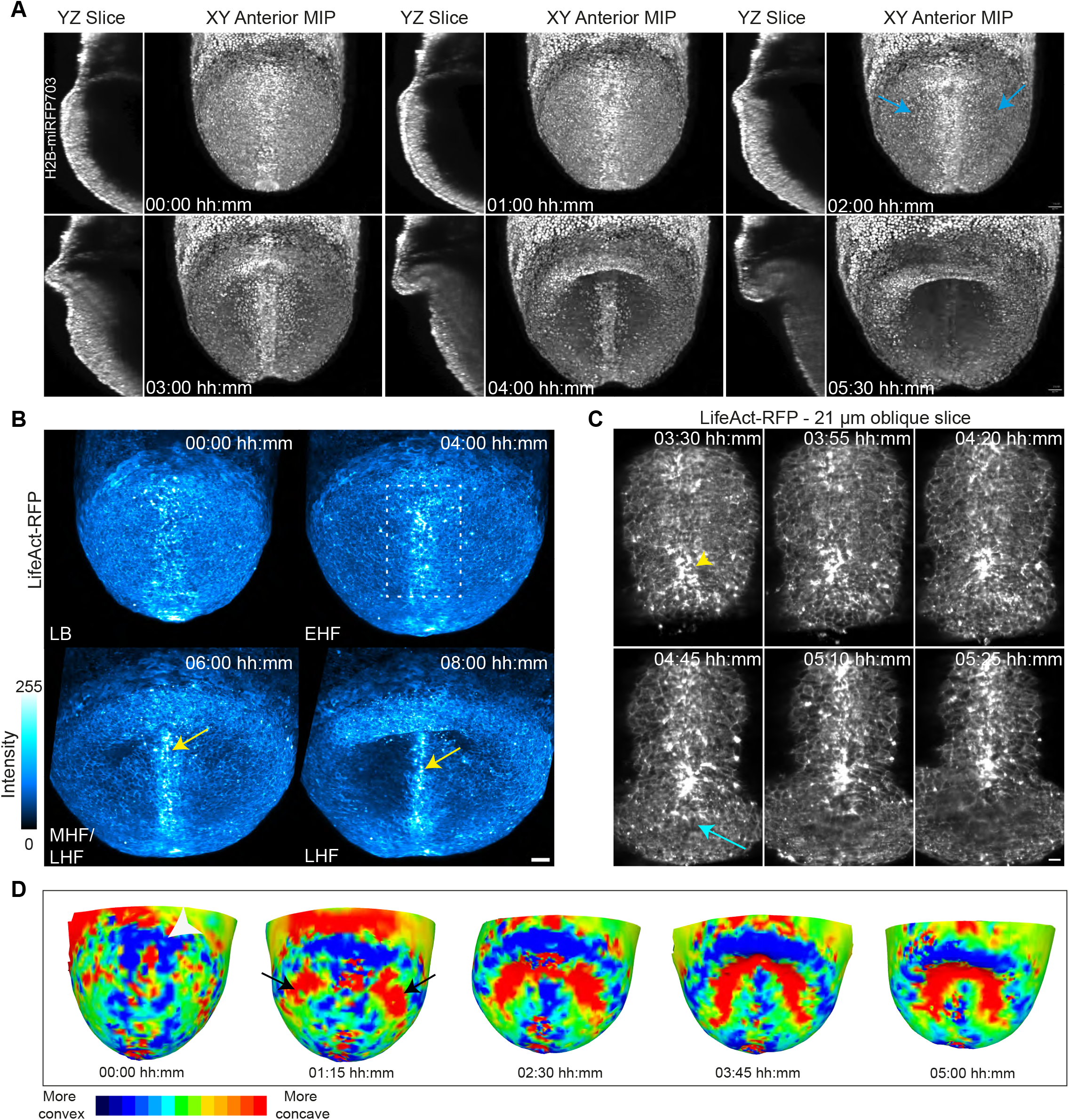
Foregut involution is a dynamic morphological process. **A** Still images from light-sheet imaging of an H2B-miRFP703 embryo from around the LB stage. MIP projections of the anterior region and 40 *µ*m lateral midline sections. Blue arrows: lateral “dimples” preceding pocket formation. Scale bar 40 *µ*m. **B** Live-imaging of a LifeAct-RFP embryo from LB to LHF stage, showing actin cytoskeleton changes over the lip of the foregut pocket and condensation over the notochord (yellow arrows). Scale bar 50 *µ*m. Outlined region expanded in **c. C** Increased actin condensation directly over the site of involution (yellow arrowhead), with inward puckering and a rapid squamous-to-columnar shape change in endodermal cells (cyan arrow). Scale bar 20 *µ*m; 21 *µ*m thick oblique slice. **D** 3D rendering of foregut involution from a simple vertex model built from live-imaging data. Curvature (quadratic fitting) is colour-mapped, with red indicating concave and blue convex regions.

Detailed examination of the morphological changes in the anterior endoderm revealed that involution proceeds through a consistent, step-wise sequence (Figure 1A). At the EB stage, the embryo possesses a rounded, cylindrical shape with a well-defined antero-posterior axis, and smooth, even surface over the entirety of the endoderm. During the late bud (LB) stage, the cardiac crescent begins to converge and a small central-proximal indentation forms at the very tip of the notochord, marking the future site of foregut involution. Notably, this indentation does not deepen further during the early headfold (EHF) stage; instead, regions on either side of the anterior notochord become concave, curving inward towards the centre of the embryo and forming distinct lateral grooves (Figure 1A, blue arrows). Only by the LHF stage does the central-proximal endoderm join these lateral indentations, involuting to produce a deeper central pocket that then elongates to form the foregut tube. Live imaging revealed that each of these stages takes approximately four hours to complete, so that the full transformation from cylindrical embryo to fully formed foregut pocket occurs within 10-12 hours (Figure 1A; Movie S1).

Live imaging of LifeAct-RFP embryos revealed regions of intense cortical actin as early as the LB stage, that were localised primarily along the central midline that were especially intense in the region of the forming foregut pocket (Figure 1B; Movie S2). This intense actin localisation along the central midline persisted throughout pocket formation, while remaining relatively lower in the lateral regions (Figure 1B). This central, linear pattern aligns with the submerging notochord, (Figure 1B, yellow arrows), making it unclear if this region of actin condensation represented a contractile structure that connected across the region of involution, or was a result of the apical surface of notochord cells condensing as they submerged (Figure 1C)^23^ and the build-up of Life-Act. The anterior tip of the notochord coincides spatially with the centre of the future foregut involution, raising the possibility that midline morphogenesis is coupled to foregut formation. The prechordal plate (PrCP) is located at the anterior tip of the notochord^20,24–26^, and therefore in direct contact with the involuting endoderm at the time of foregut formation. Although the PrCP is an essential signalling centre for anterior brain patterning, prior surgical and genetic loss-of-function studies indicate that PrCP signalling, including SHH, is dispensable for foregut tube formation *per se*^27^.

To quantify these dynamics at the tissue level, we generated three-dimensional surface meshes from point clouds created from cell nuclear positions, and computed the maximum curvature at every vertex, producing topological maps of the anterior endoderm at each developmental stage, producing a three-dimensional representation of local curvature change throughout involution (Figure 1D). This analysis confirmed that convergence of the cardiac crescent just proximal to the forming pocket precedes the involution itself, producing an outward bulge of the endoderm in the proximal region before the central foregut fully involutes at the LHF stage (Figure 1D, white arrowhead). These curvature renderings also revealed that lateral involutions consistently appear before central deepening, defining an inverted-Y geometry of the involuting endoderm at EHF (Figure 1D, black arrows).

### Morphometric analysis of mammalian embryos during ventral folding

While such vertex models provide useful three-dimensional representations of the embryo, quantitatively comparing the progression and stereotypy of foregut involution across multiple embryos requires a robust, automated, and generic morphometric approach. Such morphometric analyses would allow us to visualise tissue-scale properties, quantify changes in local tissue environments, and compare normal and mutant development directly. However, comparing the morphology of complex three-dimensional shapes that change over time is a particularly challenging computational problem, and the mouse embryo poses additional difficulties, given its hollow, cup-shaped, multi-layered structure that itself changes dramatically over the stages of interest. Additionally, embryos themselves can vary in size and shape even at the same developmental stage. We therefore sought a method capable of capturing the global shape of the embryo throughout foregut involution.

Spherical harmonics have been widely used to analyze complex shapes^28,29^, as they can reduce a complex 3D surface to a small set of coefficients that are easy to compare across samples and over time. To quantify foregut involution, we used FlowShape^22^, a spherical harmonics-based method that describes 3D shapes fully, generically and efficiently. It was previously applied to characterise cells in the *C. elegans* embryo^22^. FlowShape requires a closed triangular mesh, so we first generated meshes from light-sheet volumetric imaging of fourteen different wild-type embryos whose development covered the complete involution stages (Figure 2A). Mouse embryos are cup-shaped, and their pronounced concave curvature would dominate this curvature map, diminishing the relative contribution of the foregut folding. To avoid this, we computationally “closed the cup” (see Methods) before reconstructing the surface with an iterative shrink-wrap algorithm (Figure 2A). The method then maps mesh curvature onto a sphere using conformalised (angle-preserving) mean curvature flow, reducing the shape to a single scalar function that is decomposed into spherical harmonics.

**Figure 2:**
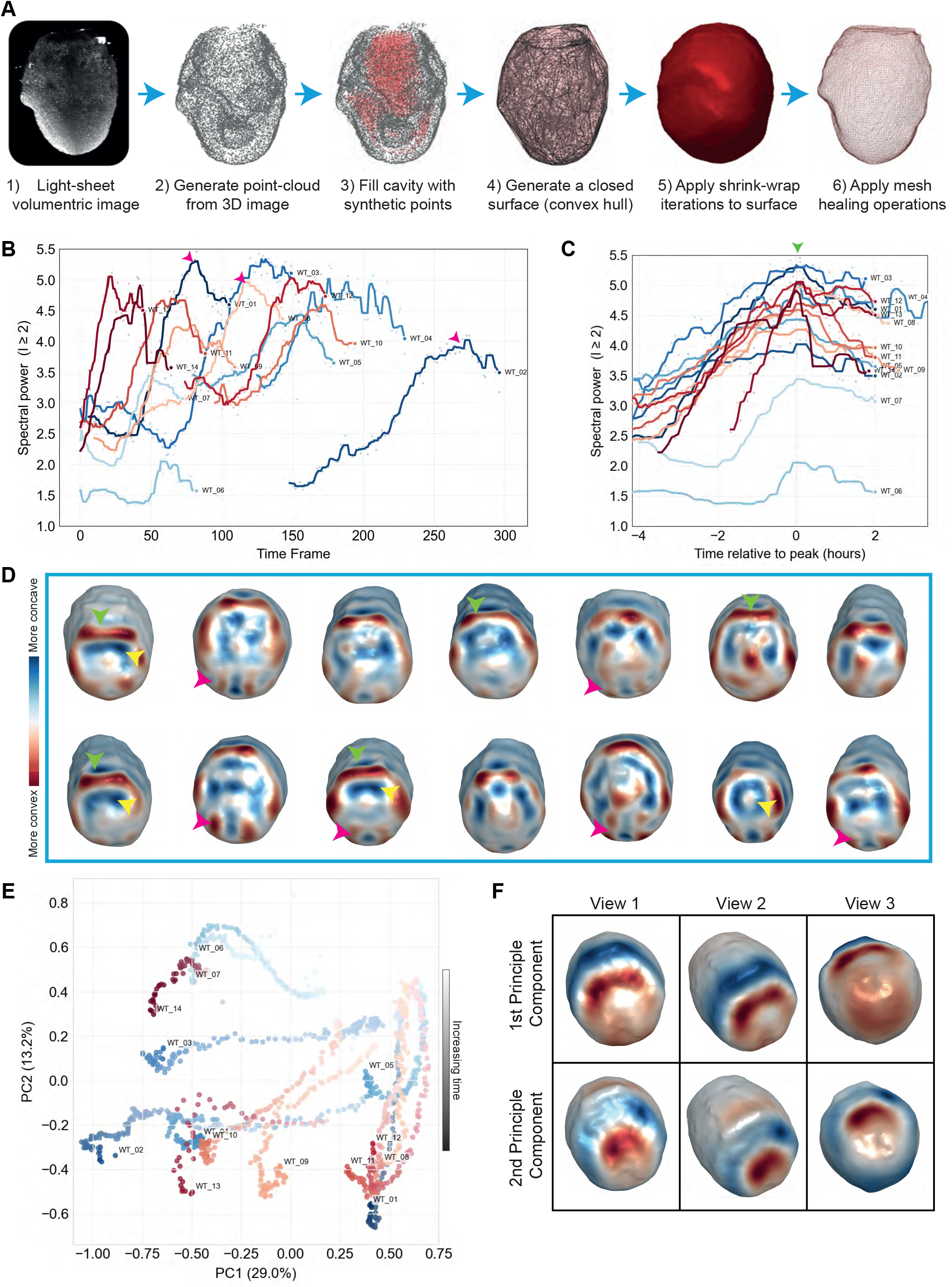
Morphometric analysis of foregut involution using spherical harmonics. **A** Computational pipeline for robust mesh generation from light-sheet data. Three-dimensional image volumes of cup-shaped mouse embryos (E7.0–E8.0) are converted into watertight surfaces, which are then decomposed into spherical harmonics using FlowShape^22^ to yield unbiased shape descriptors. **B** Shape complexity over time. Complexity is measured by the spectral power: the total energy in the spherical harmonic coefficients at each time point. The x-axis (Time Frame) shows each embryo’s own frame number from its live-imaging time series, not time normalised to a common start point; because embryos were imaged for different durations before reaching the involution window, individual traces begin at different frame numbers (e.g. WT_02 begins at frame 148, close to the onset of its own involution). Higher values indicate more shape complexity. Wild-type embryos show a stereotyped peak coinciding with maximal curvature during involution (magenta arrowheads as examples). **C** Temporal alignment of morphogenetic trajectories using the spectral powers from (B). Peak of involution is aligned to 0, green arrowhead. **D** Meshes representing the point of maximum complexity across fourteen different WT embryos. Arrow-heads highlight particular features as noted in the main text. **E** Morphospace defined by the first two principal components of the spherical harmonic descriptors, with each embryo’s developmental trajectory plotted from lighter (earlier timepoints) to darker (later timepoints) points. Trajectories cluster close together early in development, when embryos are most similar in size and shape, and progressively disperse as embryos grow in size and morphological complexity — reflecting increasing developmental variability over time and across experiments. **F** PC1 and PC2 projected on an embryo mesh. Colours indicate how PCs change with the local curvature (red, positive; blue, negative), localising the dominant modes of variation to the involuting anterior endoderm. Views 1-3 are different rotations of the same embryo.

Spherical harmonics can capture broad, simple features, such as the main axis of elongation, in a few coefficients. More complex shapes require more coefficients, to represent the finer-grained variation in mean curvature across the sphere. The sum of squared coefficients, the spectral power, captures shape complexity with a single scalar measure (see Methods). Across fourteen wild-type embryos, this power increased steadily over time, peaking at maximum involution (Figure 2B), a consistent trend which allowed us to align embryos in time (Figure 2C,D). This alignment in time highlighted several consistent physical features that are present during foregut involution (Movie S3). Namely, a pronounced ‘bulge’ just proximal to the pocket, in the region of the cardiac crescent (Figure 2D, green arrowheads); the centre of involution itself (Figure 2D, yellow arrowheads); and, lateral to the concave midline and node, ‘ridges’ or more convex features (Figure 2D, magenta arrowheads). These features are not readily apparent from volumetric imaging data, and it remains to be seen whether they reflect distinct local mechanical properties, such as stiffness.

For visual inspection of how embryo shapes change relative to each other, we performed a PCA on all 256 spherical harmonic coefficients for the WT embryos, and plotted each embryo’s time-lapse as a trajectory (Figure 2E). We visualized the most variable features by mapping PC1 and PC2 back to a shape, which confirmed that these capture the main developing morphological features, such as the involution region with the neighbouring ridge and the broader midline region (Figure 2F).

Together, these analyses show that foregut involution is a highly stereotyped process, and this unbiased spatial and temporal alignment allows for direct, quantitative comparisons between wild-type and mutant or perturbed embryos.

### The role of actomyosin in foregut involution

Having characterised foregut involution on the tissue-scale, we next sought to uncover cellular and molecular players that might be driving the involution process. The degree and localisation of actin condensation over the site of involution and the notochord suggested actomyosin contractility may be playing a role in shaping the developing pocket. However, examination by immunofluorescence revealed that phospho-myosin II was absent from the central midline (Figure S1B), indicating that this condensation of actin is not producing a contractile actin cable, although it may be providing local tissue stiffness.

To test whether actomyosin contractile forces are however required for involution, as is the case in epithelial invaginations in other organisms^30–34^, we cultured EB-LB stage embryos in the presence of the myosin II inhibitor Blebbistatin or the Rho-kinase inhibitor Rockout for up to 24 hours. Long-term Blebbistatin culture (10-25 *µ*M) compromised general embryo health (as did DMSO) but did not prevent involution: embryos (4/4 across two independent experiments) still formed an inward fold of the anterior endoderm (Figure S1C). The Rho-kinase inhibitor Rockout (50 *µ*M) was better tolerated and resulted in reduced pMLC2 staining (Figure S1D), however, all Rockout-treated embryos (8/8 across three independent experiments) initiated foregut involution and formed a recognisable foregut pocket, although the region between the pocket and the cardiac crescent was visibly malformed and adopted a pointed, rather than smooth, semicircular shape (Figure S1E, asterisk). Together, these results show that actomyosin contractility shapes some later aspects of ventral folding morphogenesis but is not required for the initial involution stages, consistent with observations in the chick foregut, where early involution is similarly driven by differential growth between the endodermal and mesodermal layers rather than by actomyosin contraction^35^.

### Anterior embryonic visceral endoderm cells undergo a spatiotemporally restricted apoptosis during foregut involution

Live imaging of *Afp*-kGFP and *Ttr*-Cre^*TG/+*^; Rosa26R^*mTmG*^ (both specific markers of visceral endoderm cells) embryos consistently revealed a striking, highly localised cell death event in the anterior endoderm just prior to and during foregut involution, which appeared to be restricted solely to the emVE cells. This event was also observed by Batki and colleagues^36^ while this manuscript was in preparation, however the timing, lineage specificity, extrusion mechanics, and morphogenetic relevance of this event have not yet been described.

Adaptive light-sheet microscopy provided simultaneous tracking of dying cells in the anterior and posterior endoderm at single-cell resolution across the entire E7.5 window (Figure 3A; Movies S4). Cell death was identified by stereotypic membrane blebbing followed by extrusion from the epithelium. Sporadic events were detectable as early as the EB stage, but the rate of cell death rose sharply from the LB stage onward, peaking concurrently with foregut involution (Figure 3A, arrows). Little cell death was observed in the posterior endoderm during this same window (Figure S2A). A small uptick in posterior emVE cell death accompanied the later onset of hindgut involution at E8.0, but cell death in the posterior endoderm remained a rare event and most posterior emVE cells were instead carried into the hindgut tube intact (Figure S2A), which may account for the higher proportion of emVE cells retained in the midgut and hindgut at E8.5^16^.

**Figure 3:**
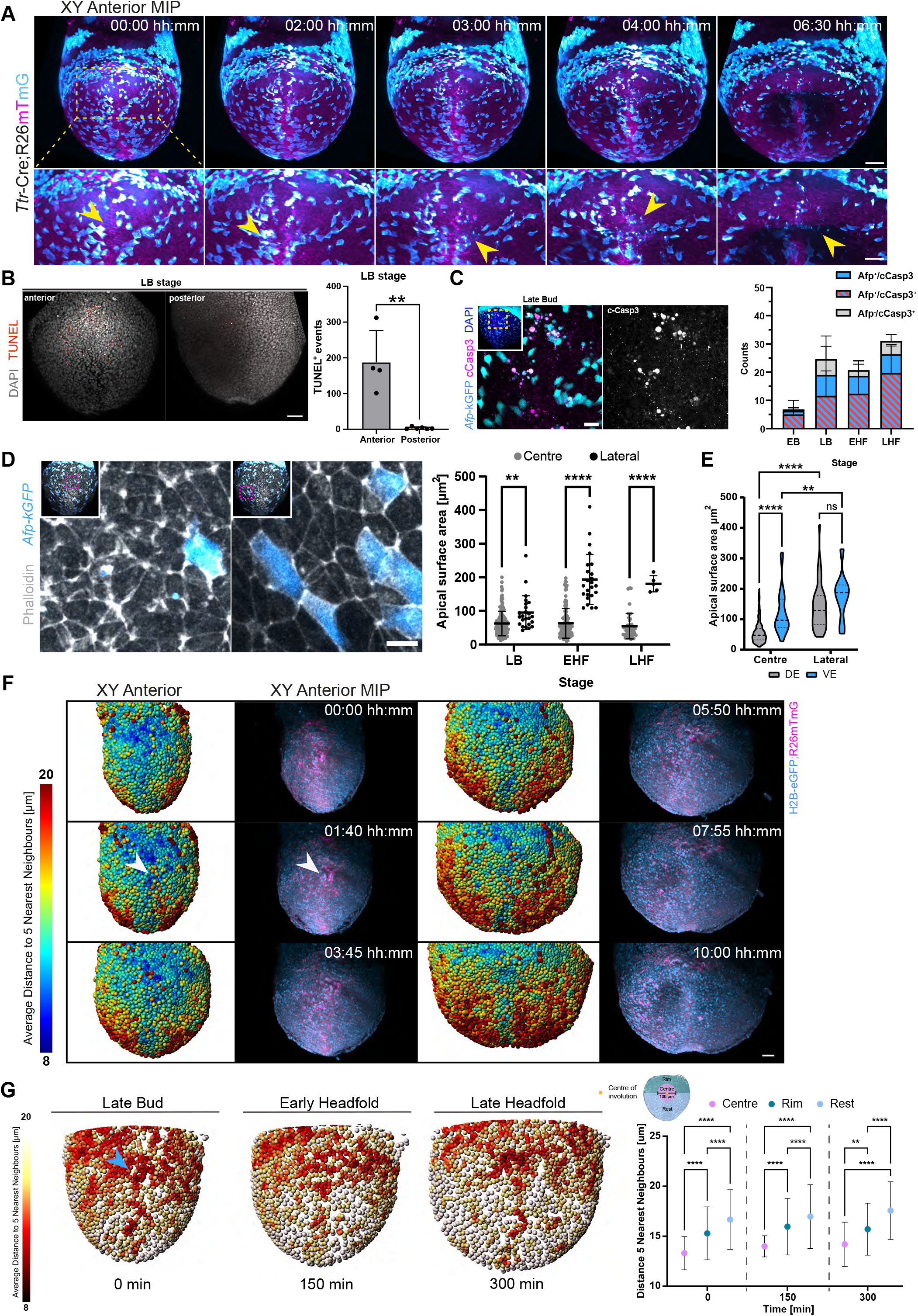
Cell death and density in the anterior embryonic visceral endoderm. **A** Selected anterior MIPs from live imaging of a *Ttr*-Cre;ROSA26^mTmG^ embryo. Yellow arrows: visceral endoderm cells undergoing cell death, indicated by blebbing and extrusion from the epithelium. Scale bars 100 *µ*m (top), 50 *µ*m (bottom). **B** (left) Anterior and posterior views of stage-matched LB embryos stained for DAPI (grey) and DNA damage (TUNEL, red). Scale bars 50 *µ*m. (right) Quantification of TUNEL-positive cells in the endoderm. N = 4 (ant.), 6 (pos.). Mann-Whitney test, ** *P* ≤ 0.01. Mean and SD. **C** (left) Close-up MIP of the forming foregut in a late bud (LB) stage embryo expressing *Afp*-kGFP (cyan), immunostained for cleaved caspase 3 (cCasp3, magenta) and DAPI (blue). Scale bar 20 *µ*m. (right) Quantification of fragmented, cCasp3-positive cells in the anterior endoderm. N = 4 (EB), 5 (LB), 3 (EHF), 3 (LHF) embryos; n = 27, 123, 87, 93 cells, respectively. Mean and SD. **D** Apical surface area of endoderm cells in the central-proximal (grey) vs. lateral (black) region across stages. LB: N = 4 embryos, n = 186 cells; EHF: N = 5, n = 144; LHF: N = 1, n = 51. Compared within each stage by Mann-Whitney test. P-values: ** ≤ 0.01, **** ≤ 0.0001. Line and error bars show mean and SD. **E** Apical surface area of emVE and DE cells (a proxy for cell density) in the central-proximal (left) vs. lateral (right) endoderm. VE cells (cyan), DE cells (grey). N = 12 embryos, n = 412 cells. Two-way ANOVA with Sidak’s multiple comparison test. P-values: n.s. *>* 0.05, ** ≤ 0.01, **** ≤ 0.0001. Scale bar 10 *µ*m. 1 pixel median filter on Phalloidin channel. **F** Cell density from selected time points of an H2B-eGFP embryo, with whole-embryo tracks and lineages from^37^. White arrow: centre of involution. The anterior-most proximal region and the midline become progressively more crowded over time (average distance to 5 nearest neighbours; blue = crowded, red = sparse). Scale bar 50 *µ*m. **G** (left) Anterior views of selected time points from a *Foxa2*-eGFP live-imaging data set. Spots are colour-coded by average distance to the 5 nearest neighbours (scale 8–20 *µ*m). (right) Quantification of cell density (distance to 5 nearest neighbours) in the anterior endoderm at three time points (0 min = LB, 150 min = EHF, 300 min = LHF stage), by region (regions depicted upper left). N = 1 embryo, n = 2649, 2536, 3014 cells at t = 0, 150, 300 min, respectively. Two-way ANOVA with Tukey’s multiple comparisons test. P-values: n.s. *>* 0.05, ** ≤ 0.01, **** ≤ 0.0001.

To quantify the degree of cell death during these stages, we performed TUNEL labelling on stage-matched LB embryos and quantified DNA fragmentation in dissected anterior versus posterior halves. TUNEL-positive nuclei were strongly enriched in the anterior half (4 anterior versus 6 posterior embryos, *p <* 0.01; Figure 3B). Quantification in *Afp*-kGFP-labelled embryos revealed that *>*80% of TUNEL-positive nuclei in the anterior endoderm were emVE in identity (Figure S2B).

Cleaved caspase-3 (cCasp3) immunostaining in *Afp*-kGFP-labelled embryos confirmed that the majority of dying cells, identified by nuclear fragmentation and blebbing, were positive for active effector caspase, consistent with classical programmed apoptosis (Figure 3C). Additionally, the majority of cCasp3-positive cells also expressed *AFP*-kGFP, confirming that this cell-death event is largely specific to the emVE. The peak of cCasp3-positive emVE cells occurred at the LB stage with progressively fewer events at EHF and LHF, mirroring the live-imaging time-course. A subset of fragmented emVE cells were cCasp3-negative; possibly a result of cell death and fragmentation before the fairly late-stage cCasp3 activation^38,39^, but also leaves open the possibility of a parallel caspase-independent route in a minority of cells.

#### Increased local cell density, but not cell proliferation, marks the future site of involution and cell death

We next asked whether localised cell behaviours within the anterior endoderm could account for the stereotypic position of emVE-specific cell death. As endoderm cells at this stage form a thin, squamous epithelium, the apical surface area of an endoderm cell can serve as a reasonable proxy for cell size and packing. Quantification of apical surface area from phalloidin-stained, *Afp*-kGFP-expressing embryos revealed that endoderm cells at the future site of involution had significantly smaller apical surfaces than cells in the lateral regions of the anterior endoderm (Figure 3D). This reduction was already present at the LB stage, before any visible inward bending, and increased through subsequent stages (Figure 3D). At the same time, apical surface area measurements showed that emVE cells in the central region had significantly larger apical surfaces than neighbouring DE cells (Figure 3E), pointing to a lineage-specific response to packing within the endoderm layer.

To determine whether this reduction in apical area reflected an increase in cell density, we used existing cell tracking results from a ubiquitously-expressing nuclear reporter^37^ as well as from live imaging and tracking (via Imaris and manual curation) of *Foxa2*-eGFP-expressing embryos, which broadly label endoderm (both definitive, and weakly VE) in addition to primitive streak, notochord, and node cells^40^. In both cases, we used Imaris to compute and visualise the average distance to the five nearest neighbours as a proxy for cell density. Cell nuclei density maps were rendered directly onto the embryo (Figure 3F, G; Movie S5). These results indicated elevated nuclear density at the central-proximal anterior endoderm even before the appearance of the central indentation. Density was consistently lower in regions lateral to the midline, including the regions that go on to undergo lateral involution at EHF (Figure 3F, G). To quantify this further, we manually divided the anterior endoderm into three distinct regions: the centre (50 *µ*m radius around the centre of involution), the proximal rim (proximal to the centre of involution

(COI), *>*50 *µ*m distance to COI) and the distal region (distal to the COI, *>*50 *µ*m distance to COI) (Figure 3G, plot). The centre region corresponds to the site of foregut invagination, while the rim region reflects the position of the forming heart field. Analysis revealed that the centre region exhibits significantly higher cell density from the early LB stage through to the full invagination of the foregut at the LHF stage, as shown by reduced distances to the five nearest neighbours (Figure 3G, plot). The increased density is most concentrated immediately anterior to the future pocket, near the tip of the notochord (Figure 3F, white arrow; Figure 3G, blue arrow). In addition, the majority of cell death events also occurred within 100 µm of the site of involution (Figure S2C,D).

To determine whether this increase in cell crowdedness was perhaps a consequence of increased local cell proliferation, we mapped pHH3-positive mitotic events in fixed wild-type embryos across the early involution stages at E7.5, just prior to the observed cell death events. pHH3-positive cells were uniformly distributed across the anterior endoderm, with no enrichment in the central region of involution at any stage (Figure S2E). Live tracking of mitoses in *Foxa2*-eGFP embryos confirmed that division events were temporally and spatially uncorrelated with the centre of involution, with rates slightly lower in the centre region than in the surrounding endoderm when normalised to local cell number (Figure S2F-I).

Together, these data argue that the localised increase in cell density at the central-proximal endoderm reflects the compression of cells into a region that does not itself proliferate at higher rates than its surroundings. A similar increase in cell density along the prospective foregut has been described in the avian embryo, where it has been proposed to generate a median hinge point that biases inward folding^41^; our data identify an analogous morphological feature in mouse and place it in temporal register with the morphological progression of foregut involution.

#### Embryonic visceral endoderm cells extrude bidirectionally through assembly of a contractile actomyosin cable

The rapid fragmentation, blebbing, and extrusion of dying emVE cells prompted us to investigate the cellular mechanics underlying their removal from the endodermal epithelium. Cell extrusion in canonical vertebrate epithelia proceeds through the assembly of a contractile, myosin-rich actin purse-string in either the dying cell or its neighbours, biasing the dying cell to the apical surface^42,43^. To test whether emVE extrusion follows this paradigm, we examined basement membrane attachment, cytoskeletal dynamics, and the direction of extrusion in live and fixed embryos.

Live imaging of *Afp*-kGFP; *LamininC1*-tdTomato embryos showed that ∼60% of dying emVE cells (52 events across 5 embryos) retained continuous attachment to the laminin basement-membrane layer in the time window immediately preceding extrusion, while ∼40% had already detached (Figure 4A, B). Cell death therefore proceeds regardless of basement membrane attachment status, ruling out anoikis as the dominant trigger.

**Figure 4:**
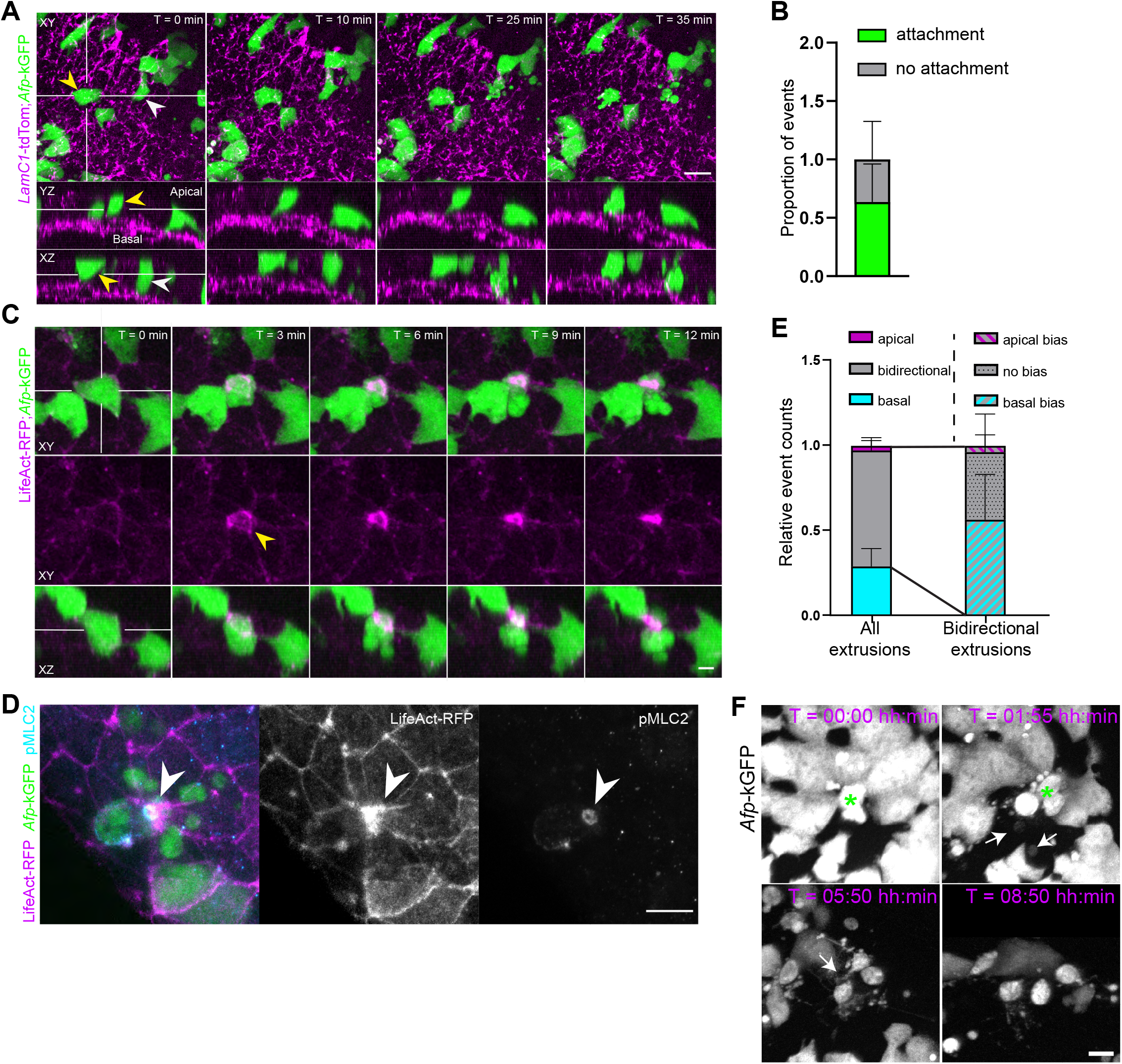
Basement membrane attachment and bidirectional extrusion in visceral endoderm cell death. **A** Selected time points of live imaging of the anterior endoderm in E7.5 embryos expressing *Afp*-kGFP (green) and *LamininC1*-tdTomato (magenta). (top) 20 *µ*m MIP. (bottom) 20 *µ*m *x*-*z* and *y* -*z* sections. Yellow and white arrowheads: two cells undergoing cell death, identified by blebbing and extrusion from the epithelium. Scale bar 5 *µ*m. **B** Quantification of basement membrane attachment in the 5 min preceding cell death. N = 5 embryos, n = 52 cells. Mean and SD. **C** Selected stills from live imaging of an E7.5 embryo expressing *Afp*-kGFP (green) and LifeAct-RFP (magenta; 3*×*3*×*1 median filter), showing emVE cell extrusion and formation of a contractile actomyosin ring. Yellow arrowhead: site of extrusion and ring formation. Scale bar 10 *µ*m. **D** Extruding emVE cell labelled by *Afp*-kGFP (green), immunostained for pMLC2 (blue) and Phalloidin (magenta; 1 pixel median filter). The actin ring is rich in pMLC2 (white arrow). Sum projection of a 20 *µ*m z-stack. Scale bar 10 *µ*m. **E** Normalised event rate relative to total quantified extrusion events. Left bar: all extrusion events. Right bar: subset with bidirectional extrusion, showing relative bias toward apical, basal, or unbiased extrusion. N = 3 embryos, n = 29 death events. Mean and SD. **F** Selected MIP stills from live imaging of apoptosis in the anterior endoderm (*Afp*-kGFP reporter). Apically extruded bodies persist for more than 8 h after cell death, forming protrusion-like structures (white arrowheads). Green asterisks: death events. Scale bar 10 *µ*m.

Live imaging using *Afp*-kGFP; LifeAct-RFP double-reporter embryos allowed us to examine the cytoskeletal behaviours surrounding this extrusion event. Just prior to fragmentation, dying emVE cells assemble a prominent F-actin ring at their apex, which then constricts (Figure 4C, yellow arrow-head; Movie S6). This ring was enriched for phosphorylated myosin light chain 2 (pMLC2) as determined by immunostaining (Figure 4D). This contractile ring did not direct extrusion in a single, apico-basally polarised manner. In *>*65% of cell death events, the actin ring constricted around the cell body without translocating the dying cell fully to either surface, producing partial bidirectional extrusion with apoptotic bodies released to both the apical and basal sides of the epithelium (Figure 4E). Among the bidirectional events, more than half were asymmetric, with greater cellular content extruded basally. Of 29 extrusions across 3 embryos, only one was unidirectionally apical; basal-only extrusion was observed in ∼28% of events. Basally extruded fragments were eventually lost to imaging, whereas apically extruded fragments persisted for hours and formed bleb-like structures on the surface of the embryo (Figure 4F, white arrows).

These observations describe a cell extrusion mode that involves a contractile actomyosin cable; basement membrane attachment that is permissive but not required; and bidirectional or basal release of apoptotic bodies.

#### Suppression of cell death in the embryonic visceral endoderm does not prevent foregut involution but results in aberrant cell aggregation

To assess whether emVE cell death is required for foregut involution, we adopted complementary chemical and genetic strategies. *Ex vivo* culture of EB-stage embryos with the pan-caspase inhibitor Z-VAD-FMK (50 or 100 *µ*M) reduced TUNEL signal in the anterior endoderm by ∼85% relative to DMSO controls (Figure 5A,B; Figure S2J). Notably, however, DMSO itself is not an inert control - even low concentrations affected embryo health. Extended Z-VAD-FMK culture for *>*21 hours blocked apoptosis throughout foregut formation, yet 10/10 embryos across three independent experiments completed foregut pocket formation (Figure S2J). Consistent with the known requirement for caspase activity in heart and neural tube morphogenesis^44^ (in addition to the effect of DMSO), embryos cultured under these conditions exhibited general developmental delays and defects in those tissues; foregut involution itself, however, was unaffected.

**Figure 5:**
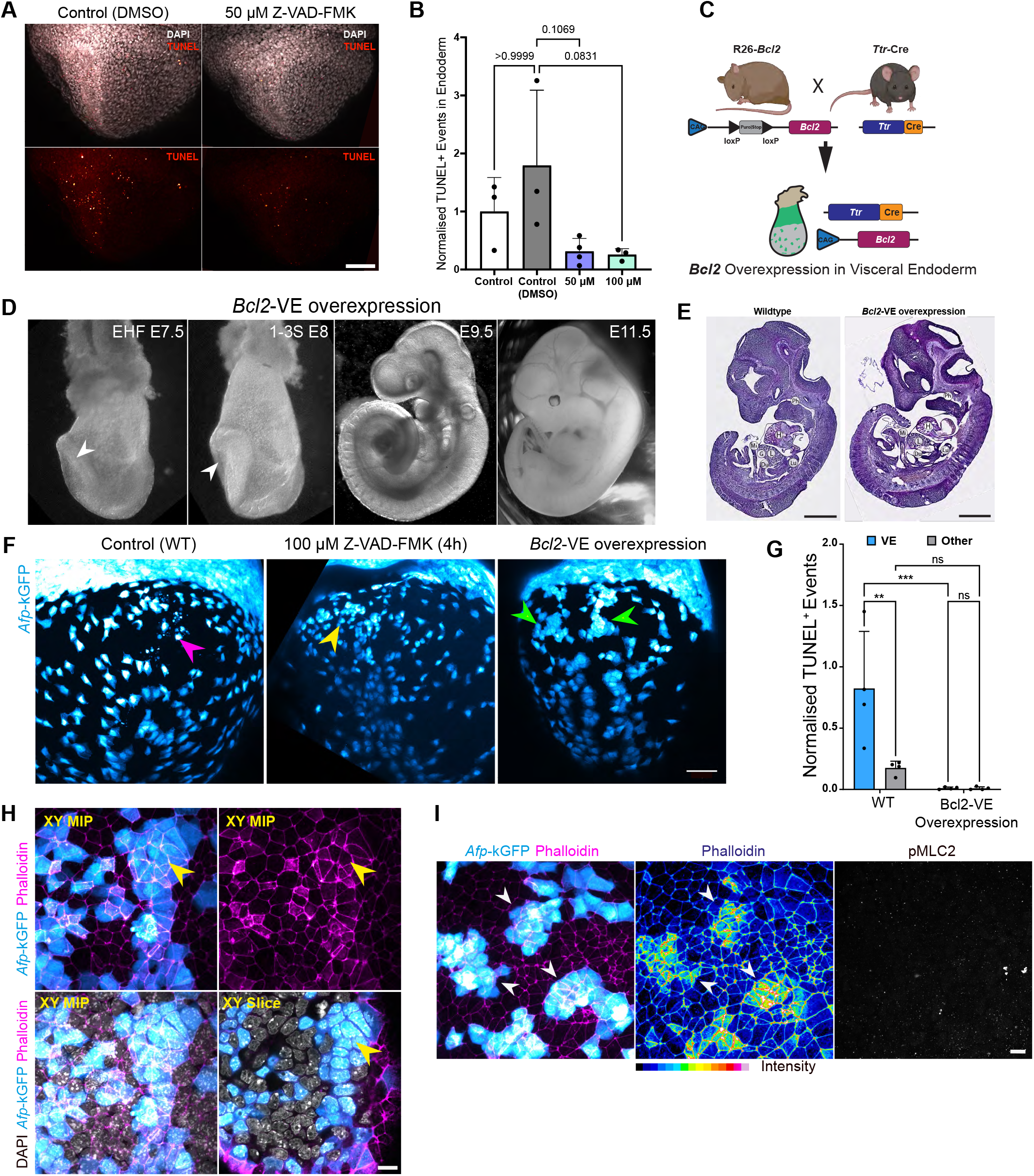
*Bcl2* over-expression in the emVE blocks cell death and causes cell aggregation. **A** Stage-matched EHF embryos cultured *ex vivo* for 4 h in DMSO or 50 *µ*M pan-caspase inhibitor (Z-VAD-FMK), stained with DAPI (grey) and TUNEL (red). Scale bar 100 *µ*m. **B** Quantification of TUNEL-positive events in the anterior endoderm after 4 h culture *±* Z-VAD-FMK, normalised to control medium without DMSO. N = 3 (Control, Control DMSO, 100 µM Z-VAD-FMK), 4 (50 µM Z-VAD-FMK) embryos, n = 354 (Control), 635 (Control DMSO), 76 (50 µM Z-VAD-FMK), 91 (100 µM Z-VAD-FMK) events. Kruskal-Wallis test with Dunn’s multiple comparison test. Mean and SD. **C** Schematic of crossing scheme to achieve *Bcl2*-overexpression specifically in VE cells using a *Ttr*-Cre. **D** Brightfield images of E7.5–E11.5 embryos with VE-specific *Bcl2* overexpression. Headfolds elevate by E7.5 and the anterior foregut pocket is fully formed by E8.0 (white arrowheads). Embryos survive to at least E11.5 with no obvious phenotypic abnormalities versus wildtype. **E** Haematoxylin & Eosin staining of an E11.5 wildtype and *Bcl2*-VE overexpressing embryo. H: Heart; Ph: Pharynx; Lu: Lung bud; L: liver primordium; G: gall bladder primordium; Du: duodenum; MI: midgut. Scale bar 1 mm. **F** Representative MIP images of LB stage embryos expressing the VE cell marker *Afp*-kGFP. VE cells form aggregates in the absence of cell death in (*Bcl2*-VE overexpression but not in 4-hour culture in 100 *µ*M pan-caspase inhibitor Z-VAD-FMK, 1 pixel median filter). Arrowheads point to region (or expected region) of emVE cell death. Scale bar 50 *µ*m. **G** Quantification of TUNEL-positive events of the anterior endoderm of WT and *Bcl2*-VE overexpressing embryos. N = 4 (WT, all LB stage), 4 (*Bcl2*-VE overexpression, 3 LB stage, 1 EHF stage) embryos, n = 750 (WT), 15 (*Bcl2*-VE overexpression) TUNEL-positive events. Two-way ANOVA with uncorrected Fisher’s LSD. P-values: ** *P* ≤ 0.01, *** *P* ≤ 0.001. Mean and SD. **H** Close-up of a VE cell aggregate in a *Bcl2*-VE overexpressing embryo. Scale bars 10 *µ*m. 1 pixel median filter applied in Afp (cyan) and Phalloidin (magenta) channels. **I** Immunostaining for F-actin (Phalloidin, magenta and intensity heat-map) and pMLC2 in *Bcl2*-VE overexpressing embryo expressing *Afp*-kGFP. Cell aggregates are rich in apical actin (arrowheads, Phalloidin intensity heat-map) but show no presence of activated myosin. 1 pixel median filter applied. Scale bar 10 *µ*m.

To restrict the loss of cell death specifically to emVE, we crossed mice expressing *Ttr*-Cre, which drives recombination in embryonic and extraembryonic VE,^45^, with a conditional Rosa26-*Bcl2* allele to overexpress the anti-apoptotic factor BCL2^46,47^ in emVE (Figure 5C; *Bcl2*-VE). *Bcl2*-VE embryos showed a complete loss of TUNEL-positive cells in the anterior endoderm at E7.5 (Figure 5G), confirming the specificity and efficiency of the manipulation. Despite this, *Bcl2*-VE embryos completed foregut pocket formation, established the foregut tube, and proceeded through ventral folding and turning (Figure 5D). H&E staining at E11.5 revealed normal embryo morphologies, and recognisable lung, liver, gallbladder, and gut derivatives were all present (Figure 5E). The foregut and its derivatives therefore form correctly in the absence of emVE cell death.

To determine whether emVE death contributes more subtly to involution dynamics, we examined *Bcl2*-VE embryos co-expressing the *Afp*-kGFP reporter (Figure 5F; Movie S7). Strikingly, in the absence of emVE death, the normally sparse emVE cells failed to disperse and instead formed multilayered aggregates across the entire anterior endoderm (Figure 5F, green arrowheads). This wide-spread aggregation was not seen in either control or Z-VAD-FMK treated embryos (Figure 5F, pink and yellow arrowheads denoting site where cell death is or would occur). These aggregates contained cells with elevated cortical F-actin, and occasionally included multinucleated cells and cells with abnormal nuclear morphology (Figure 5H, I). Both the aggregation phenotype and increase in cortical F-actin is consistent with the non-canonical role of *Bcl2* in mediating cell-cell contact inhibition^48,49^. This suggests that emVE cells maintain their largely salt-and-pepper distribution across the endoderm layer via cell-cell contact mediated inhibition, and the increased cell-crowdedness experienced by emVE cells over the site of involution triggers apoptosis in these cells, which can be blocked by over-expression of *Bcl2*.

### Lineage- and stage-resolved bulk RNA-seq distinguishes anterior emVE, anterior DE, and their posterior counterparts

The surprising result that actomyosin contractility was not required for foregut involution coupled with a lineage-specific cell death event led us to ask if signalling or other molecular factors were involved in or over-represented during these stages. Existing RNA-sequencing data^1,50–52^ does not cover the precise stages of foregut involution with the temporal resolution needed, and as emVE cells are only a fraction of the total cell population in the embryo there is a risk that they are under-represented. Additionally there may be differences between anterior and posterior regions of the embryo at these stages. We therefore generated a new RNA-sequencing dataset which discriminates between emVE and DE, both posterior and anterior, over precise stages during foregut involution.

To uncover the molecular signatures of emVE and DE cells during the short period of foregut involution, we used *Afp*-kGFP; *Foxa2*-eGFP doubly-transgenic embryos at four different and precisely collected stages (EB, LB, EHF, LHF). Embryos were dissected into anterior and posterior halves, dissociated into single-cell suspension, and sorted by FACS into emVE (kGFP^+^/eGFP^+^), DE (kGFP^−^/eGFP^+^), and Mes/Epi (kGFP^−^/eGFP^−^) populations (Figure 6A-B). kGFP is a variant of GFP, and is a single, possibly random, mutation away from Folding Reporter GFP (a precursor to superfolder GFP)^53,54^. Significant differences in intensity between the *Afp*-kGFP and *Foxa2*-eGFP reporters allowed for a clean separation of the three populations by FACS (Figure 6B). 112 embryos were collected (34 EB, 30 LB, 27 EHF, 21 LHF) and sequenced as biological triplicates per stage, position, and population (Table S1).

**Figure 6:**
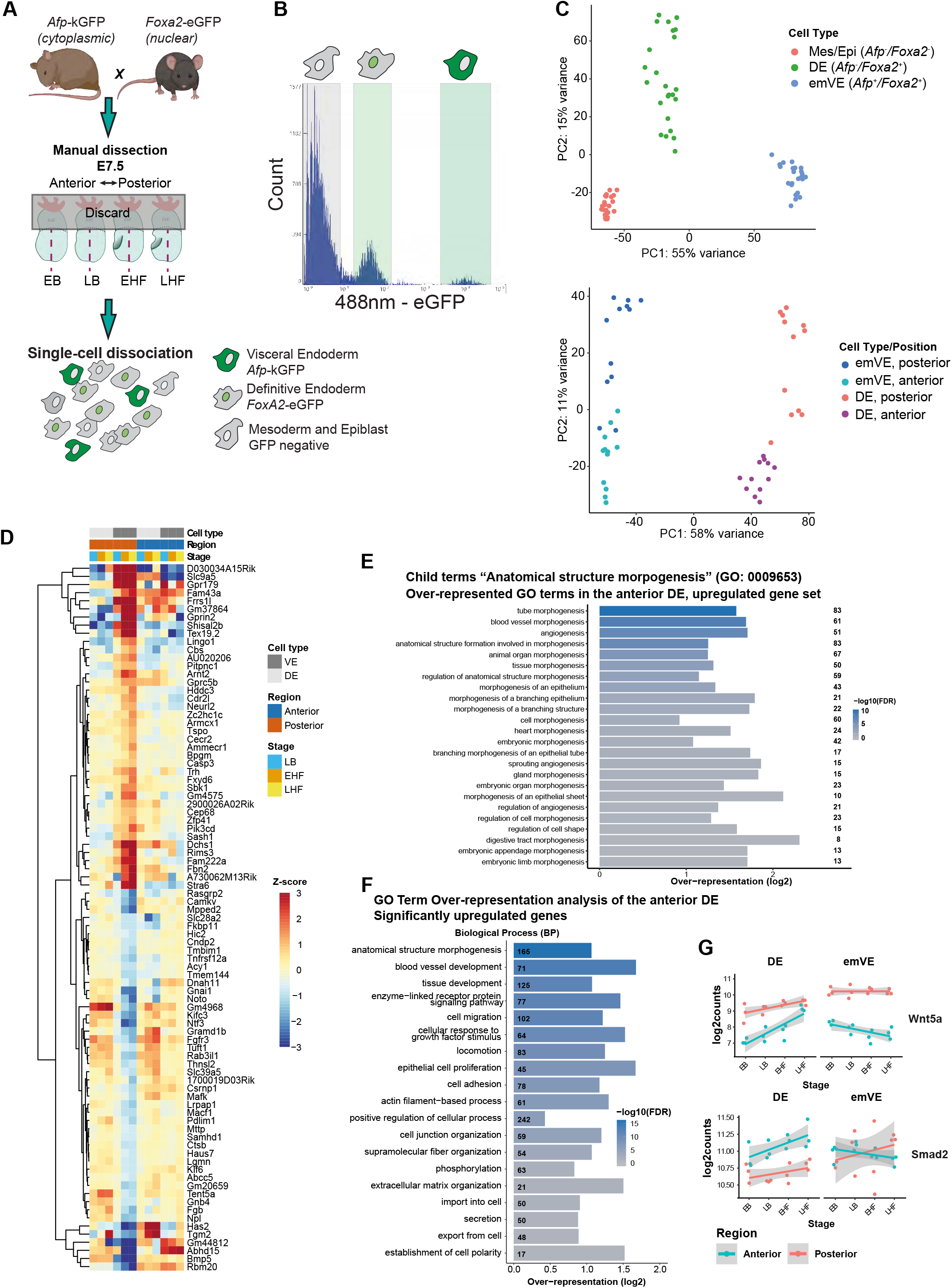
Bulk RNA sequencing of spatial and temporally staged embryos. **A** Sample collection strategy for bulk RNA-seq. Simultaneous labelling of emVE (*Afp*-kGFP) and DE (*Foxa2*-eGFP) cells enables endoderm enrichment. **B** Representative FACS profile of dissociated embryos. eGFP and kGFP intensities are discriminable based on intensity, allowing separation of emVE and DE cells. **C** Principal component analysis (PCA) of transcriptional differences based on the top 5,000 most variable genes. (Top plot) Mesoderm/epiblast (Mes/Epi), definitive endoderm (DE) and embryonic visceral endoderm (emVE) samples. (Bottom plot) DE and emVE samples only; PC1 separates samples by cell type, PC2 by position. **D** Heatmap of 87 of the 186 genes whose abundance changed across E7.5 specifically in posterior emVE — significant in posterior emVE and in interaction tests against anterior emVE and posterior DE (see Methods) — restricted to genes with absolute log_2_ fold-change per stage > 0.25. Values are normalised to the EB-stage mean within each lineage × position group **E** Over-represented child terms of GO:0009653, “anatomical structure morphogenesis”, among the significantly overexpressed genes in the anterior DE relative to the posterior DE and anterior emVE. Adjusted *P <* 0.01. Numbers to the right indicate the number of differentially expressed genes belonging to each GO term. **F** Over-represented GO terms among significantly upregulated genes in the anterior DE vs. anterior emVE and posterior DE, by Biological Process (BP). **G** Expression of *Wnt5a* and *Smad2* in the anterior DE and emVE, shown as separate plots for each gene. Dots represent individual replicates; lines show the fitted linear regression with confidence interval.

Principal component analysis recovered a clean, hierarchical separation of cell identity, position, and stage (Figure 6C, D). PC1 separated emVE from DE, PC2 separated anterior from posterior endoderm, and within-population clustering ordered samples by developmental stage (Figure 6C, top and bottom plots). Pairwise comparisons confirmed that DE diverges more strongly across stages than emVE, consistent with the rapid morphogenetic specification of gastrointestinal organ progenitors in the DE during this window (Figure S3A-G). For all subsequent analyses we modelled stage as a continuous variable and used interaction terms with cell type and position to identify lineage-, position-, and stage-specific gene expression changes (Figure S3A-G; Methods).

#### Anterior DE cells specifically upregulate organ morphogenesis and adhesion programmes

Differential expression analysis identified 1,118 genes whose abundance changed across the E7.5 sub-stages differently in anterior DE than in anterior emVE (567 up, 551 down). The upregulated set was strongly enriched for GO terms governing organ morphogenesis (Figure 6E). 83 of 567 (∼15%) upregulated genes annotated to “tube morphogenesis” (GO:0035239), with multiple child terms covering heart and vascular development as well as “digestive tract morphogenesis” (Figure S3I). Genes downregulated in anterior DE were enriched for cilium-related GO terms, consistent with the displacement of the Foxa2+, ciliated node toward the posterior across the staging window (Figure S3J).

Cell adhesion (GO:0007155) was a top-enriched parent term among genes upregulated in anterior DE, encompassing 78 of 567 (∼14%) upregulated genes (Figure 6F, Figure S4A). Reactome pathway analysis highlighted enrichment of both cell-cell and cell-matrix adhesion programmes, including multiple integrin-, collagen-, and laminin-mediated pathways (Figure S4B). Five candidate adhesion regulators emerged with distinct expression patterns (Figure S4C). Cadherin-3 (*Cdh3*; P-cadherin) and cadherin-6 (*Cdh6*; K-cadherin) were more abundant in anterior emVE than in anterior DE at the earliest sub-stages, but rose steeply across E7.5 specifically in the anterior DE (Cdh6, +1.29 log_2_ per stage, adj. P = 7 × 10^−25^; Cdh3, +0.49 log_2_ per stage, adj. P = 1 × 10^−11^), converging on emVE levels by EHF. The actin-binding protein *Lima1* (EPLIN), a linker between the cadherin-catenin complex and F-actin, was most abundant in anterior emVE, where it declined across E7.5 in a manner specific to that population (-0.25 log_2_ per stage, adj. P = 7 × 10^−15^). Conversely, junctional adhesion molecule 3 (*Jam3*) and nectin-1 (*Nectin1*) were least abundant in anterior emVE. Nectin1 additionally showed an inverse position-specific trajectory, declining in anterior emVE while increasing in posterior emVE across E7.5 (Figure S4C). The combination of elevated *Lima1* and reduced *Nectin1* in anterior emVE provides a candidate molecular basis for the differential cell-cell coupling, accumulation of cortical actin in surviving cell aggregates, and altered (bidirectional) extrusion behaviour we observe in this population.

#### Anterior emVE specifically downregulates cell cycle and chromosome segregation programmes

Anterior emVE cells showed few specific transcriptomic changes - 144 genes (65 up, 79 down) relative to anterior DE and posterior emVE - and no GO term enrichment among upregulated genes. Downregulated genes, however, were enriched for cell-cycle terms (GO:0022402, “cell cycle process”; 20 of 79 genes, 2.6-fold, adj. P = 6.0 × 10^−3^), with the strongest enrichment for the more specific term “sister chromatid segregation” (GO:0000819; 12 genes, 6.9-fold, adj. P = 4.7 × 10^−4^) (Figure S3K, S5A). 20 of the 79 downregulated genes were annotated for cell cycle regulation, including the kinesin motors *Kif15* and *Kif2c* and the kinetochore component *Ndc80*. Reactome analysis reinforced enrichment for kinetochore function and sister-chromatid segregation. Crucially, this down-regulation was specific to anterior emVE: posterior emVE and both anterior and posterior DE retained normal expression of these genes (Figure S5B, C).

Posterior emVE cells, by contrast, were defined by 186 differentially expressed genes (77 up, 109 down) relative to anterior emVE and posterior DE, and yielded only a single over-represented term (“vesicle”) among the downregulated genes and none among the upregulated (Figure 6D). Comparable in size and effect magnitude to its anterior counterpart, the posterior set nonetheless converges on no common programme. The two populations differ less in how much their transcriptomes change than in how concertedly they change: only in the anterior emVE does that change resolve into a single coherent programme, the coordinated withdrawal of mitotic machinery, coincident in space and time with the crowding and apoptosis described above.

#### Differential expression of anti- and pro-apoptotic genes in the anterior DE

Transcriptomic analysis of genes associated with programmed cell death (GO:0012501 and GO:0043069) revealed two significantly upregulated genes in this category (*Foxp1* and *Bex2*) and two significantly downregulated anti-apoptotic genes (*Rtkn2* and *Arg2*) in anterior emVE cells compared to anterior DE and posterior emVE cells. Notably, both *Foxp1* and *Bex2* have been reported to suppress apoptosis in other cell types, and in the case of *Bex2* through upregulation of *Bcl2*^55,56^. This raises the possibility that their induction in the anterior emVE reflects an intrinsic but ultimately insufficient pro-survival response in this population. This would be consistent with our finding that a stronger, exogenously-imposed anti-apoptotic signal (*Bcl2* overexpression) is sufficient to fully block cell death in the same population, whereas the endogenous response evidently is not.

While anterior DE cells exhibited a broader set of differentially expressed genes relative to the anterior emVE cells associated with apoptosis regulation, none particularly stood out in terms of known function or expression dynamics (Figure S6A,B) aside from *Ccng1*, (Figure S6A). As a member of the G-type cyclins, *Ccng1* has been implicated in modulating cellular sensitivity to apoptotic stimuli and increased expression of *Ccng1* is associated with heightened susceptibility to radiation-induced cell death^57,58^. *Ccng1* overexpression can promote activation of *Ccnb1* (Cyclin B1), facilitating exit from G2 arrest and subsequent cell death, in a manner that is thought to be P53-independent^57^. Similar to the effects of elevated P53 levels in MDCK cells^59^, elevated *Ccng1* does not induce cell death in an unperturbed state of the cell but rather makes cells more sensitive to apoptosis upon perturbation^58^. Whether the increased expression of *Ccng1* in emVE cells translates to an appreciable increase in the protein level remains to be tested. However, it is tempting to speculate that the mechanical stresses acting on the emVE cells during foregut formation, caused by the increased crowdedness and invagination of the tissue, might trigger apoptosis in a cyclin G1 and cyclin B1 dependent way, similar to that observed during exposure to radiation in fibroblasts and lung cells in culture^57,58^.

#### A receptor and Smad expression landscape consistent with an emVE-selective BMP2 response

Madabhushi and Lacy previously demonstrated that VE-specific deletion of *Bmp2* produces a disorganised anterior phenotype with mispositioned headfolds and heart and absent or aberrantly placed foregut^5,10^. Examination of the bulk RNA-seq dataset showed that *Bmp2* expression at E7.5 is highest in emVE, with significantly higher transcript levels in anterior emVE than posterior (Figure S7B), consistent with previous *in situ* hybridisation results from Lacy and colleagues^10^. Among the digestive-tract-morphogenesis genes, *Smad2* and *Wnt5a* showed differential expression, with *Smad2* enriched in anterior DE and *Wnt5a* strongly enriched in the posterior endoderm of both lineages (Figure 6G).

Among type I BMP receptors, *Bmpr1a* was uniformly expressed across all populations and stages, *Bmpr1b* was expressed at consistently low levels, lowest in emVE, and *Acvr1* expression was strongly enriched in emVE (Figure S7C). Type II receptors *Bmpr2, Acvr2a*, and *Acvr2b* were all expressed; emVE cells displayed slightly higher *Bmpr2* and lower *Acvr2b* than DE and Mes/Epi (Figure S7D). Among canonical Smads, *Smad2, Smad3*, and *Smad5* were broadly expressed, with *Smad5* declining in emVE through E7.5; inhibitory Smads *Smad6* and *Smad7* and the BMP-specific *Smad9* were uniformly low (Figure S7E-F). This expression landscape positions VE-derived BMP2 as a candidate ligand acting through an *Acvr1*/*Bmpr2*-rich emVE compartment in addition to a more uniformly *Bmpr1a*-driven epiblast compartment, supporting the genetic observation that mosaic *Bmpr1a* deletion in the epiblast phenocopies several features of *Bmp2*-VE^60^ and identifying *Acvr1* and *Smad5* as the most plausible mediators of BMP2’s anterior-specific role.

### VE-specific deletion of *Bmp2* links midline morphogenesis to foregut involution

To revisit this mutant in the context of the cellular and molecular framework established above, we crossed *Bmp2*^flox/flox^ mice with *Ttr*-Cre to ablate *Bmp2* specifically in the VE (*Bmp2*-VE KO; Figure 7A). Consistent with the original description, *Bmp2*-VE KO embryos at E8.0 showed pronounced anterior disorganisation, with twisted, buckled headfolds and a misplaced foregut pocket (Figure 7B, C, Figure S7A). Notably however, the posterior development of these embryos was strikingly normal. The node progressively retracted over the distal tip towards the posterior, and somite formation, tailbud elongation and hindgut formation proceeded as in wild-type embryos. In total, 5 of 6 *Bmp2*-VE KO embryos across three independent litters managed to form a shallow foregut pocket or indentation (Figure 7C), albeit in an aberrant position and with delayed timing, indicating at least the initial process of foregut involution remained intact in these mutants, which was not evident in the previous study. This may reflect background-related differences (CD1 in this study versus C57BL/6 in^10^) and suggests that BMP2 signalling from the VE is playing an indirect role in directing foregut involution.

**Figure 7:**
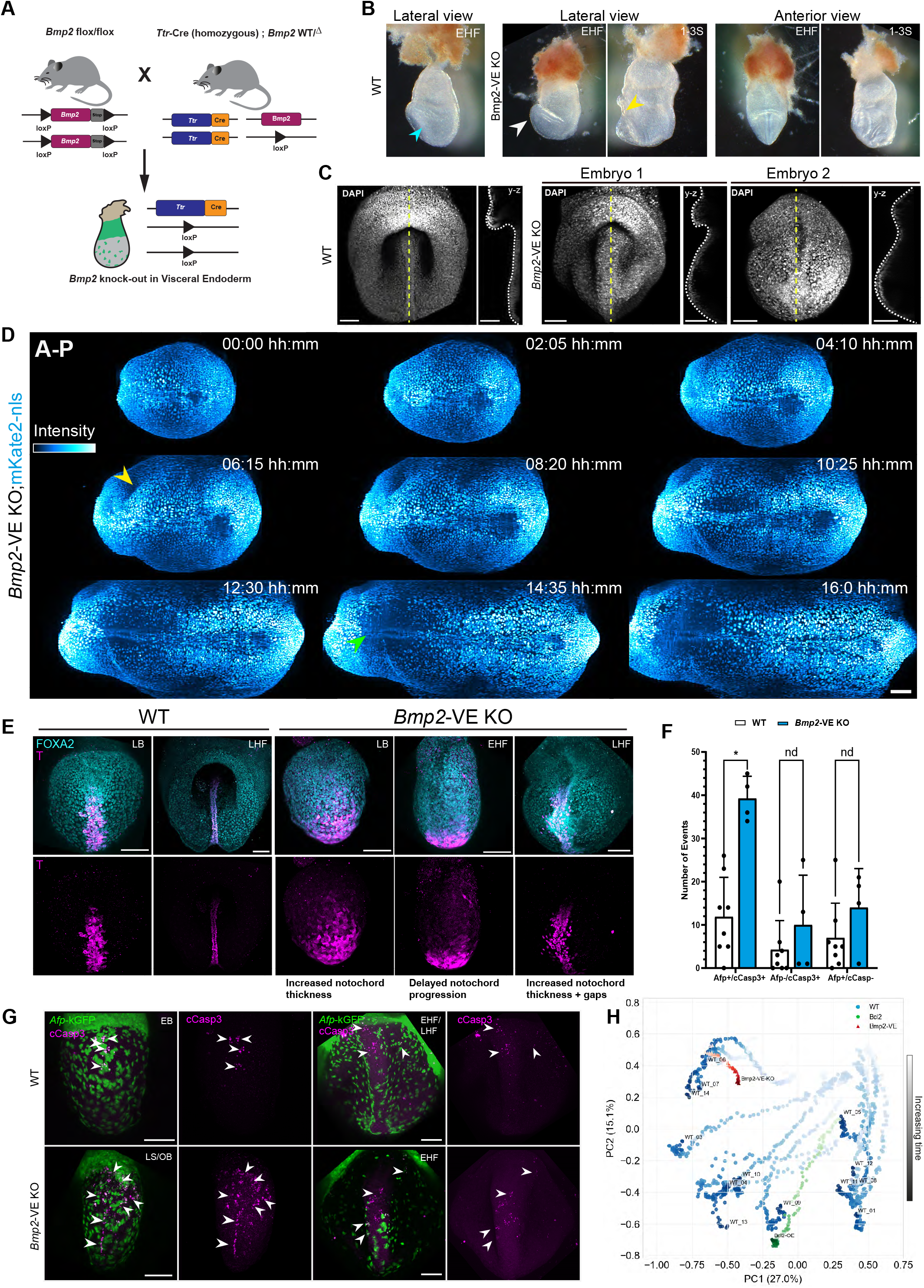
VE-specific *Bmp2* ablation disrupts anterior development and foregut positioning. **A** Crossing scheme for VE-specific *Bmp2* knockout. **B** Brightfield images of *Bmp2*-VE KO embryos at EHF and 1–3 somite stages. **C** MIPs of a WT and two LHF stage *Bmp2*-VE KO embryos stained with DAPI, with orthogonal *z*-*y* slice views at the yellow dashed line (white dashed line: endoderm layer). Scale bar 100 *µ*m. **D** Light-sheet imaging of a *Bmp2*-VE KO embryo expressing mKate2-nls (cyan; anterior left, posterior right, YZ Projection). Headfolds begin to buckle (yellow arrowhead), and no pocket formation is visible, while cardiac contraction onset (green arrowhead, line artefacts) and posterior development proceed normally. Wavelet fusion, 50 *µ*m size 1 pixel median filters. Scale bar 100 *µ*m. **E** Anterior views of wildtype and *Bmp2*-VE KO embryos immunostained for Foxa2 (cyan) and Brachyury (magenta). MIPs. Scale bars 100 *µ*m. **F** Quantification of cCasp3-positive death events in the anterior endoderm of fixed WT and *Bmp2*-VE KO samples (LB and EHF stages). VE cells are *Afp*^*+*^, DE cells *Afp*^−^. N = 8 (WT), 4 (*Bmp2*-VE KO) embryos. Multiple Mann-Whitney tests with Benjamini, Krieger and Yekutieli FDR correction (5% FDR). P-values: * *P* ≤ 0.05. **G** Increased cell death in *Bmp2*-VE KO embryos. Immunostaining for cCasp3 (magenta) in *Afp*-kGFP (green) WT and *Bmp2*-VE KO embryos from EB to EHF/LHF stages. Arrowheads: cCasp3-positive and/or fragmented cells in the endoderm layer. cCasp3 channel despeckled, with outliers removed (5 pixel radius). Scale bars 100 *µ*m. **H** Morphospace represented by the first two principal components of the spherical harmonic descriptors. Wild-type embryos trace a reproducible trajectory defining normal involution, dispersion captures experimental variation. Visceral-endoderm-specific BCL2 overexpression follows a trajectory indistinguishable from wild type, whereas *Bmp2-VE KO* produces an aberrant trajectory.

Live imaging of *Bmp2*-VE KO embryos expressing mKate2-nls showed that ventral folding attempts to proceed in mutants but loses its normal central symmetry, with aberrant folding and buckling of the neural headfolds (Figure 7D, yellow arrowhead; Movie S8). The position of the expanding headfolds correlated tightly with the position of lateral endoderm involutions in the same mutant embryos (Figure 7D, yellow arrowhead), implying mechanical coupling between the elevating headfolds and the endoderm. In embryos that failed to form a central foregut pocket, proper positioning of the heart-field was also perturbed, however the onset of cardiac contractions began at the expected developmental time (Figure 7D, green arrowhead). These linked deformations place ventral folding into a single mechanical unit.

In addition to the dramatic morphological defects in the heart, headfold, and foregut we observed that the notochord often appeared disrupted in mutant embryos. To examine whether the anterior disorganisation in *Bmp2*-VE KO embryos is related to defects in the underlying axial mid-line, we immunostained mutants and littermate controls for Brachyury and Foxa2 (markers of the notochord and endoderm and notochord, respectively) across E7.5 stages. In wild-type LB embryos, the notochord is 5 to 6 cells wide and extends from the node approximately halfway to the proximal anterior border; by LHF it has narrowed to ∼2 cells across, and its anterior most tip extends to the centre of the future foregut pocket (Figure 7E). In *Bmp2*-VE KO embryos, notochord morphogenesis was severely perturbed: at LB the notochord was either overly broad or its proximal extension was delayed; by LHF it remained wider than wild-type and frequently displayed gaps that broke its continuity (Figure 7E). These defects are similar (but do not exactly phenocopy) midline phenotypes seen in epiblast-specific *Lhx1* mutants in which VE displacement is disturbed and node/midline morphogenesis fails^61^, and *FoxH1* deletion mutants^62^. While the notochord’s role in patterning the neural tube via signalling is well established^27,63–66^, these results suggest that the early notochord may also play a structural role in anterior morphogenesis, providing a rigid substrate against which the elevating neural headfolds can “push” upward. Consistent with this, when the notochord is absent or disorganised — as in *Bmp2*-VE KO embryos — the neural headfolds continue to expand, proliferate, and even bend, but fail to elevate properly, producing the “bulging” anterior morphology and variable phenotypes seen across embryos.

#### Spatial restriction of emVE apoptosis is lost in *Bmp2*-VE KO embryos

Cell death in the anterior endoderm of *Bmp2*-VE KO embryos was no longer spatially confined. cCasp3 immunostaining of stage-matched *Afp*-kGFP; *Bmp2*-VE KO embryos showed a clear increase in apoptotic cells across the anterior endoderm and an earlier onset relative to wild-type, with elevated emVE cell death already evident at the late streak/early-bud stage - well before the wild-type peak at LB (Figure 7F,G). The defining spatial confinement of emVE death to the central proximal foregut field, the hallmark of the wild-type phenotype, was lost in the mutant; cell death occurred broadly across the anterior endoderm, in addition to being elevated (Figure 7G).

#### Morphospace analyses of mutant vs wild-type developmental trajectories

Using FlowShape to align and map the developmental trajectories of developing embryos over time, we next included both *Bcl2*-VE over-expression and *Bmp2*-VE KO embryos into our morphometric analyses. While movies obtained from light-sheet imaging of *Bcl2*-VE embryos visually appeared to have initially shallower and slower-forming foregut pockets, plotting the morphogenetic trajectory using spherical harmonics alongside wild-type embryos did not produce a convincingly different curve (Figure S8A, B, Figure 7H, green trajectory). In contrast, when we projected a *Bmp2*-VE mutant into the same space, it followed a markedly aberrant trajectory and lacked the peak in power over time that characterises involution. (Figure 7H, red trajectory, Figure S8C).

## Discussion

The work presented here provides the first integrative account of the cellular, mechanical, and molecular determinants of foregut involution in the mouse embryo, and places this event within the wider context of ventral folding morphogenesis. Three findings in particular are central to this account. First, foregut involution proceeds through a stereotyped morphological progression and does not require actomyosin contractility for its initiation. Second, a spatiotemporally restricted wave of apoptosis, triggered by cell-crowding in a sensitive population, removes a precisely defined cohort of anterior emVE cells through a contractile actin ring; while this death is ultimately dispensable for involution, emVE cells show pronounced aggregation when cell death is inhibited via upregulation of *Bcl2*. Third, anterior VE-derived BMP2 regulates foregut involution indirectly, by guiding notochord morphogenesis and the global mechanics of the headfolds and heart field, rather than by triggering the onset of involution.

The differential adhesion landscape uncovered by lineage-resolved RNA-seq, elevated *Lima1* and reduced *Nectin1* and *Jam3* in anterior emVE, with parallel upregulation of *Cdh3, Cdh6*, and integrin-mediated adhesion programmes in anterior DE, provides a candidate molecular basis for both the selective extrusion of anterior emVE and their aggregation when extrusion is blocked. Functional testing of these candidates, in combination with the cell-cycle/chromosome-segregation programme that is selectively downregulated in anterior emVE, will be required to determine which molecular features are causally responsible for the unique behaviours of this population. The dataset itself, with stage-, position-, and lineage-resolution across the involution window, provides a resource for hypothesis generation across many additional aspects of anterior morphogenesis.

The study here also presents several remaining questions. The mechanical coupling between elevating headfolds and the proper positioning of the notochord, suggested by the tight spatial correlation between headfold positions and axial midline in *Bmp2*-VE KO embryos, will require targeted laser ablation or direct force inference to confirm. The down-stream signalling axis through which VE-derived BMP2 directs the notochord to the anterior invites comparative RNA sequence profiling between wild-type and *Bmp2*-VE KO embryos. And the ontogeny of the cell-cycle and adhesion programmes that distinguish anterior from posterior emVE remains to be traced back to their AVE origins. Resolving these questions will require continued advances in live imaging, mechanical perturbation, and lineage-resolved transcriptomics, building on the framework established here.

In order to place the process of foregut involution in a quantitative framework we utilised spherical harmonics to find the peak of involution and align embryos in an unbiased manner in space and time. By generating a representative 3D system we are able to project multi-modal information onto a single framework, which allows for the cross-comparison of different metrics such as curvature and cell-direction or velocity. While the *Bmp2*-VE KO mutant produces a severe morphological phenotype that is readily identified by our morphometric analyses, this method could also be used to identify more subtle changes in embryo shape and development in less severe mutants or to compare across developmental stages.

This careful dissection of foregut involution in the mouse embryo has revealed it to be a model process for studying morphogenesis in which lineage-specific cellular behaviours, tissue-scale mechanics, and spatially restricted signalling converge on a single shape change. The dissection of this event reframes ventral folding as a coordinated geometric problem - heart, headfolds, foregut, and midline arranged around a shared template - rather than as a sequence of independent organogenic events, and provides additional understanding for how the anterior body plan is mechanically and molecularly assembled.

## Acknowledgements

JK, GSN, SAG, HW, LK, PS, TS, and KM are supported by the Medical Research Council as part of UK Research and Innovation (MCUP1201/23). RJ is supported by an FWO research grant G008423N and KU Leuven internal funds grants C14/24/109 and IDN/25/004. CvB is supported by FWO aspirant grant 11L0923N. For the purpose of open access, the MRC Laboratory of Molecular Biology has applied a CC BY public copyright licence to any Author Accepted Manuscript version arising. We thank the LMB FACS facility, in particular Pier Andree Penttila for their help in preparing the bulk RNA seq dataset, the LMB Biomed team and ARES for all animal work and husbandry, the LMB mechanical and electrical workshop, and the LMB light-microscopy facility. Gerry Crossan and Pranay Shah developed the *Bcl2* over-expression line, and kindly allowed us to use and describe it in this study. We thank members of the LMB Cell Biology division for their helpful comments on this manuscript.

## Data availability

All data supporting the findings of this article are available from the corresponding authors on reasonable request. Bulk RNA sequencing data used in this manuscript will be uploaded to a public repository associated with this manuscript. Code and software used for morphometrics analyses and image processing can be found in Methods.

## Methods

### Experimental model and subject details

#### Mice

All animal experiments performed in this study were approved by the Medical Research Council’s Laboratory of Molecular Biology animal welfare and ethical review body and conform to the UK Home Office Animal (Scientific Procedures) Act 1986 (Licence no. PP5976836). Mice were maintained under specific pathogen-free conditions in independently ventilated cages (GM500; Techniplast) on Lignocel grade 2/2 or FS-14 spruce bedding (IPS) at 19-23 °C. All mice were kept on a mixed C57BL/6J;129Sv;CD1 or CD1 background.

#### List of mouse strains

The following mouse lines were used in this study: *Afp*-kGFP (Tg(Afp-GFP*)#Maba/J; kind gift from K. Hadjantonakis^67^); *LamininC1*-tdTomato^68^; *Foxa2*-eGFP (RIKEN;^69^); *Bmp2*^flox/flox^ (B6;129S4-Bmp2^tm1Jfm^/J, The Jackson Laboratory; Strain #016230); *Ttr*-Cre (kind gift from K. Hadjantonakis; MGI:3829595)^45^; Rosa26R^mTmG^ (STOCK Gt(ROSA)26Sor^tm4(ACTB-tdTomato,-EGFP)Luo^/J; The Jackson Laboratory; Strain #007576); LifeAct-RFP (Tg(CAG-mRuby); MGI:4831038)^70^; Rosa26-*Bcl2* (kind gift from G. Crossan; see below); R26-CAG-nuc-3xmKate2-nls (RIKEN)^71^; and wild-type mice on a mixed or CD1/Hsd:ICR background (Inotiv; MGI:5652530).

#### Generation of the Rosa26-*Bcl2* mouse line

The Rosa26-*Bcl2* allele was a kind gift from Pranay Shah and Gerry Crossan, generated by CRISPR/Cas9-mediated knock-in at the *Rosa26* locus, following the strategy of^72^ (Full generation and validation details are described in Shah’s PhD thesis^73^). Briefly, the targeting vector encoded a CAG promoter followed by a loxP-flanked puromycin-resistance/stop cassette and the *Bcl2* coding sequence, flanked by homology arms complementary to the *Rosa26* locus on either side of the sgRNA target site. The targeting vector plasmid, sp-Cas9 and sgRNA against Rosa26 were microinjected into one-cell mouse zygotes. Injected zygotes were transferred into surrogate mothers to generate chimaeras that were screened for the presence of the transgene using endpoint PCR. Chimaeras were backcrossed with wildtype C57BL6/J mice to generate heterozygotes. Validation of the *Bcl2* over-expression can be found in^73^.

### Method details

#### Embryo collection and culture

Female mice used in timed mating experiments were aged between 8-45 weeks. Timed matings were performed overnight, and female mice were assessed for the presence of a copulation plug the following morning. The morning of the copulation plug was designated as E0.5. Females were ultrasounded at E6.5 or later to confirm pregnancy. Pregnant mice were sacrificed using cervical dislocation. The uterus was transferred into a polystyrene petri dish (Corning, cat. no. 353003) with dissection medium on a heating plate (okolab, H401-Nikon-SMZ-180-glass) at 37 °C. Embryos were recovered by manual dissection from the decidua using forceps, followed by the removal of the Reichert’s Membrane using tungsten needles. Dissection of embryos was performed in dissection medium (DMEM FluoroBrite (Gibco, A1896701) supplemented with 1% Penicillin Streptomycin, 1% MEM NEAA (100x) (Gibco, 11140-035), 1% GlutaMAX-I (100x) (Gibco, 35050-061) and 10% heat-inactivated Fetal Bovine Serum (FBS) (Corning, 35-016-CV)) for all live imaging and culture experiments or phosphate-buffered saline (PBS) supplemented with 10% qualified FBS (Fisher Scientific Ltd, 11560636) for experiments on fixed samples. Samples were immediately used or fixed in 4% paraformaldehyde (in PBS) for further analysis.

Embryos were cultured in 50% rat serum (in-house production or Janvier Labs) in DMEM FluoroBrite (Gibco, A1896701) supplemented with 1% Penicillin Streptomycin, 1% MEM NEAA (100x) (Gibco, 11140-035), 1% GlutaMAX-I (100x) (Gibco, 35050-061) and 10% heat-inactivated Fetal Bovine Serum (FBS) (Corning, 35-016-CV). Short-term culture for up to 6 hours was performed in Ibidi 8 well slides (Ibidi, cat. no. 80806) in 400 *µ*L culture medium. Long-term culture (*>*6 hours) was performed in 24-well plates (Corning, cat. no. 3526) in 500 *µ*L culture medium on an orbital shaker (Sanyo) at 120 rpm. Up to 2 embryos were placed per well. Culturing was performed at 37 °C and 5% CO_2_.

Inhibitors supplemented to the culture medium for perturbation experiments were the pan-caspase inhibitor Z-VAD-FMK (Millipore, cat. no. 627610), (*±*)-Blebbistatin (ChemCruz, cat. no. sc-203532), and the Rho-kinase inhibitor Rockout (ChemCruz, cat. no. sc-203237). Final concentrations used for experiments can be found in the corresponding results sections and figure legends. For drugs dissolved in DMSO (Sigma Aldrich, cat. no. D2438), controls were performed with the medium supplemented with the corresponding concentration of DMSO as well as in culture medium without any supplementation to control for the effect of DMSO itself.

#### Immunofluorescence staining and imaging

Dissected embryos were rinsed in PBS and fixed in 4% paraformaldehyde (in PBS) for 1 h on ice for *Afp*-kGFP-expressing strains, and 4 °C over-night while rocking for all others. Subsequently, blocking was performed in 10% Donkey Serum (Abcam, cat. no. AB7475) in PBS-T (0.1% Triton) overnight at 4 °C. Primary antibodies were diluted in blocking solution, and incubation was performed for 1-4 nights at 4 °C while being placed on a shaker. After washing of samples in blocking solution for 30-45 minutes at room temperature, incubation with secondary antibodies diluted in blocking solution was performed overnight at 4 °C or for 4 h at room temperature. Antibodies and their concentrations used in this study were: anti-Foxa2 (Abcam, cat. no. ab108422; 1:200), anti-Brachyury/T (R&D Systems, cat. no. AF2085; 1:50), anti-cleaved caspase-3 (Asp175) (5A1E) (Cell Signaling Technology, cat. no. 9664; 1:200), anti-phospho-Histone H3 (Ser10) (Sigma-Aldrich, cat. no. 06-570; 1:400), and antiphospho-myosin light chain 2 (Thr18/Ser19) (Cell Signaling Technology, cat. no. 3674; 1:200). Secondary antibodies, all used at 1:1000, were Alexa Fluor 405-conjugated donkey anti-rabbit (Invitrogen, cat. no. A48259), Alexa Fluor 488-conjugated donkey anti-rabbit (Invitrogen, cat. no. A21202), Alexa Fluor 555-conjugated donkey anti-goat (Invitrogen, cat. no. A27039), Alexa Fluor 555-conjugated donkey anti-rabbit (Invitrogen, cat. no. A31572), and Alexa Fluor 647-conjugated donkey anti-rabbit (Invitrogen, cat. no. A32795). Nuclei were counterstained with DAPI (ThermoFisher Scientific, cat. no. 62248; 1:1000), and F-actin was visualised with Alexa Fluor 405- or Alexa Fluor 647-conjugated Phalloidin (Invitrogen, cat. nos. A30104 and A22287, respectively; 1:250). After a washing step in PBS for 30 minutes, samples were immediately used for imaging.

For confocal imaging of fixed samples, embryos were placed in 15 Well 3D Glass Bottom slides (Ibidi, cat. no. 81507) and embedded in 0.5% low-melting-point agarose (Biogene, 300-800) in PBS. Samples were oriented with the anterior side facing the coverslip using microloader gel loading tips (Eppendorf, cat. no. 10289651).

Images were taken on a Zeiss 710 or 780 inverted confocal microscope, using a 20x/0.8NA air or 63x/1.4NA oil objective with a z-stack thickness of 2 *µ*m for images acquired at 20x magnification and 1 *µ*m for images taken at 63x magnification.

After dissection, E9.5 and E11.5 *Afp*-kGFP-expressing wild-type and *Bcl2*-VE overexpressing embryos were rinsed in PBS, fixed in 4% paraformaldehyde (in PBS) for 1 hour on ice, and then taken through a sucrose gradient (15%, 30%, and 30% sucrose:OCT 1:1) before subsequent embedding in OCT mounting media (VWR Chemicals, cat. no. 361603E). Embedded samples were sectioned to a thickness of 15 *µ*m using a cryostat (Leica CM1950) and captured on adhesion microscope slides (Epredia, cat. no. J1800AMNZ). Samples were stored after embedding or after sectioning at −70 °C until performing the immunostaining protocol.

Before immunostaining, slides were thawed at room temperature for 15 min and then rinsed with distilled water. Samples on slides were surrounded using a Hydrophobic Barrier Pen (Vector Laboratories, cat. no. VEC-H-4000). Subsequently, blocking was performed for 30 min at room temperature in 10% Normal Donkey serum (Abcam, cat. no. AB7475) in PBS-T (0.1% Triton). Primary antibodies were incubated for 1 hour at room temperature. After 2*×*15 minutes washing in PBS, incubation with secondary antibodies was performed for 30 min at room temperature. Samples were mounted in ProLong Antifade mounting medium (Invitrogen, cat. no. P36970). Primary and secondary antibodies were as listed above.

#### Haematoxylin and Eosin staining

Embedding and sectioning of samples was performed as earlier described. Subsequently, haematoxylin and eosin staining (H&E Kit, Abcam, cat. no. ab245880) was performed following the manufacturer’s instructions. After staining, slides were mounted in DPX mounting medium (Sigma-Aldrich, cat. no. 06522).

Slides (immunostaining and H&E staining) were imaged using an Olympus VS200 slide scanning microscope with a 2x/0.06NA air objective for overview scans and a 10x/0.4NA air or 20x/0.8NA air objective for detail scans.

#### Live imaging using spinning disk confocal microscopy

Embryos were dissected from the uterus as previously described, using dissection media. Up to four E7.5 mouse embryos were mounted into an imaging chamber specifically designed for this study and cultured in 400 *µ*L culture medium 50% rat serum (Janvier Labs or in-house production) in DMEM FluoroBrite supplemented with 1% Penicillin Streptomycin, 1% MEM NEAA (100x), 1% GlutaMAX-I (100x)).

For customised imaging chambers, PDMS holding moulds containing four wells (1.5 mm *×* 3 mm) were generated by cutting double-sided tape (AR-90880, 1.5 mm thickness; Adhesive Research) using a Graphtec CE6000 cutting plotter. Two of those moulds were placed into 8 well glass-bottom chamber slides (Ibidi, cat. no. 80807). Embryos were held in place with their anterior side facing the coverslip by tucking the ectoplacental cone beneath a thin glass capillary, which was attached to the PDMS moulds using dental wax (Dental Aesthetics).

Live imaging was performed using a Nikon W1 Spinning Disk Confocal with a 25x/1.05NA silicone objective. Samples were imaged every 2 to 5 minutes for up to 8 hours with a z-stack thickness of 2 *µ*m and a total imaging depth of up to 200 *µ*m.

#### Live imaging using light-sheet microscopy

Embryos were dissected from the uterus as previously described, using DMEM FluoroBrite dissection medium and 5% FBS. Whole mouse embryos were imaged on a SiMView light sheet microscope as described in^21^. Post-processing of acquired images was performed according to previously described methods^21,74^.

### RNA bulk sequencing

#### Isolation of definitive and visceral endoderm cells using fluorescence-activated cell sorting (FACS)

Simultaneous labelling of definitive and visceral endoderm cells was achieved by crossing *Foxa2*-eGFP homozygous with *Afp*-kGFP heterozygous mice. After retrieval, embryos were screened for GFP. Only embryos positive for both fluorophores were used. Embryos were staged according to Downs and Davies^75^ and embryos of the same stage (Early Bud, Late Bud, Early Headfold and Late Headfold) were pooled. Using a micro knife (Surgical Specialities Sharpoint, 72-2201), the extra-embryonic part of the embryo was cut off and discarded. Subsequently, the anterior embryonic portion was separated from the posterior half by dissection. Anterior and posterior halves of the embryo were subsequently processed separately. Samples were carefully placed in 1.5 mL Eppendorf tubes, rinsed with PBS and stored on ice until dissociation.

A single-cell suspension was generated by dissociation. 500 *µ*L Accumax (Sigma, cat. no. A7089) *+* 0.1% DNAse (Sigma, cat. no. 04716728001) was added to the samples. Samples were inverted 20*×* and then placed at 37 °C while shaking at 300 rpm for 5 minutes. After repeating this step one more time, samples were carefully pipetted once using a 1 mL tip and placed back at 37 °C (shaking, 300 rpm) for 5 minutes. Subsequently, samples were gently pipetted 10*×* using a 1 mL tip and placed again at 37 °C (shaking, 300 rpm) for 5 minutes. The single-cell suspension was filtered into BSA-coated 1.5 mL RNAse-free tubes (Invitrogen, cat. no. AM12400) using a 70 *µ*m cell strainer (Corning, 352350). For inactivation, 100 *µ*L FBS were added to the sample. Pelleting was performed at 300 g for 5 min in a centrifuge at 4 °C. The supernatant was then carefully removed, and the cells were resuspended in 200 *µ*L 10% FBS in PBS. Samples were immediately stored on ice until sorting.

Fluorescence-based cell sorting was performed using the ThermoFisher Bigfoot with a 70 *µ*m nozzle at 60 psi. The sample differential was maintained at 0.6 psi. The sample and collection device temperature was kept at 4 °C. FSC/SSC gating was used to exclude debris and doublets, and populations were defined by negative, mid and high expression levels of GFP with autofluorescence background measured using an eGFP negative control. Fluorescence was detected using a 125 mW 488 nm excitation and 507/19 bandpass filter for emission. Cells were sorted into 75 *µ*L Buffer RLT (RNeasy Micro Kit, Qiagen, cat. no. 74004) *+* 1% 14.3 M *β*-mercaptoethanol (Sigma, cat. no. M3148-100ML) and immediately homogenised by vortexing for 1 minute. Samples were stored at −70 °C until further processing.

#### RNA isolation and quality control

Before RNA isolation, samples of different sorts were pooled to achieve similar cell numbers in replicates. RNA was isolated using the RNeasy micro kit (Qiagen, cat. no. 74004) following the manufacturer’s protocol. The extracted RNA was dissolved in 14 *µ*L RNAse-free water, of which 1 *µ*L was used for RNA quality control. RNA quantification and quality control were performed using the Bioanalyzer RNA 6000 pico assay (Agilent, cat. no. 5067-1513) following the manufacturer’s protocol. All samples used for sequencing had a RIN score *>*8.0.

#### Library preparation, RNA bulk sequencing and data processing

Library preparation and RNA sequencing were performed by the CRUK-CI Genomics Core Sequencing Service. Library preparation was performed using the NEBNext Multiplex Oligos 96 Unique Dual Indexes Kit (New England Biolabs, cat. no. E6440). Sequencing was performed using the NovaSeqX, with 20M reads per sample. The pooled library balance was controlled using MiSeq nano with 1 lane of 25B reads. All sample types were run as pooled biological triplicates as earlier described and processed simultaneously.

Raw reads were processed using the nf-core/rnaseq pipeline, implemented in Nextflow and built on the nf-core framework, using the star_salmon route. Read quality was assessed with FastQC^76^ and summarised across samples with MultiQC^77^. Adapter and quality trimming was performed with Trim Galore^78^. Trimmed reads were aligned to the *Mus musculus* GRCm39 primary assembly with STAR^79^, and transcript abundances were estimated with Salmon^80^ from the resulting transcriptome-coordinate alignments. The transcriptome was generated by the pipeline from the GRCm39 primary assembly and the Ensembl release 108 annotation, and therefore includes all annotated biotypes. Transcript-level estimates were summarised to gene level with tximport, yielding counts for 56,980 genes.

#### Differential expression analysis

Gene-level counts were analysed in R using DESeq2^81^. Developmental stage was encoded in two ways: as an ordered factor (EB, LB, EHF, LHF), tested as successive pairwise contrasts, and as a continuous variable (1–4), testing for a monotonic change in abundance across the four sub-stages. The two encodings were compared directly (Figure S3A–C); the continuous encoding recovered substantially more differentially expressed genes, and genes detected only under the ordinal encoding were predominantly of very low abundance. The continuous encoding was therefore used for all subsequent analyses. Fold-changes were shrunken using lfcShrink with the ashr estimator. For analyses of change across stage within a single population, genes with an adjusted p-value < 0.01 were considered significant.

#### Definition of lineage- and position-specific gene sets

To identify changes in abundance restricted to one lineage in one position, results were combined across a main-effects model and one or two interaction models (Figure S3D), requiring both significance in every constituent model and a consistent combination of coefficient signs. Genes specific to the anterior emVE were required to change across stage in anterior emVE (main effect), and to show significant stage × cell-type (emVE vs DE, anterior) and stage × position (anterior vs posterior, emVE) interactions, with all three coefficients of the same sign. Genes specific to the posterior emVE were required to change in posterior emVE, with a stage × cell-type interaction (emVE vs DE, posterior) of the same sign and a stage × position interaction of the opposite sign, and additionally not to change significantly in the same direction in anterior emVE. Genes specific to the anterior DE were required to change across stage in anterior DE and to show a stage × cell-type interaction (emVE vs DE, anterior) of opposite sign; because the biologically relevant contrast at the site of involution is between anterior DE and anterior emVE, this set was not additionally required to differ from posterior DE. Genes specific to the anterior mesoderm/epiblast were required to change in anterior mesoderm/epiblast with a concordant stage × position interaction. Because a gene must pass the threshold in two or three independent models, a relaxed per-model threshold of adjusted p < 0.1 was used; the resulting false discovery rate for the intersection is substantially below 10%.

#### Over-representation analysis

ORA was performed with the ‘GOseq’ package^82^ using the mean read count as bias factor. GO annotations were retrieved from Ensembl (v113) via ‘biomaRt’ and expanded so that each gene was annotated to all ancestor terms of its directly annotated terms. GO terms with too few genes to afford reasonable power were removed using a procedure adapted from the DESeq2 independent-filtering approach, applied to gene-set size rather than read count. Over-represented terms were called at an adjusted *p*-value < 0.05 for the lineage- and position-specific gene sets, and < 0.01 for analyses of change across stage within a single population. Redundant terms were then removed by ranking terms by p-value and, starting from the smallest, discarding all ancestor and offspring terms with larger *p*-values, repeating until all terms were accounted for. Reactome pathway annotations were obtained from Ensembl (v113) and mapped to pathway names using ‘reactome.db’ ; the same procedure was applied, omitting the ancestor-expansion step.

### Image analysis

#### Quantification of cell death and division events

Identification and labelling of cell death and division events in live imaging data sets were done manually using Imaris. To identify cell death events, embryos expressing *Ttr*-Cre^TG/+^;ROSA26^mTmG^ or *Afp*-kGFP were used. Cell death was identified by cell fragmentation and blebbing. Tracking of cell division events in the endoderm was performed using data sets of *Foxa2*-eGFP-expressing embryos. Every cell visibly undergoing cell death or division in the next time frame, respectively, was labelled as a ‘spot’. A reference point was generated at the centre of invagination for normalisation. *x*-, *y* - and *z*-coordinates were exported as a .csv file and used for further analysis.

For the calculation of the distribution of death and division events relative to the centre of the foregut invagination, Cartesian coordinates were transformed into polar coordinates using the cart2pol function in Python, and angular and radial coordinates relative to the centre of invagination were plotted as rose plots.

All cell division events were binned according to their distance from a hand-labelled “centre of involution” reference point, and their position relative to a pair of reference points marking the proximal (boundary between embryonic and extra-embryonic tissue) and distal (node) tips of the embryo. To assign points to the “Centre” bin, we calculated the vector distance from the point of interest to the centre of involution reference point. For points beyond a distance threshold of 50 *µ*m (100 *µ*m diameter), we classified them according to the angle between the vector from the centre of involution to the point of interest and the vector from the distal reference point to the proximal reference point. If this angle was *<*90 °, the point of interest was assigned to the Rim region; otherwise, it was assigned to the Rest region.

For fixed samples, cell death was assessed by quantifying DNA fragmentation using terminal deoxynucleotidyl transferase dUTP nick-end labelling (TUNEL), performed with the Click-iT Plus TUNEL assay (Invitrogen, cat. no. C10618) following the manufacturer’s protocol. To analyse divisions in fixed samples, immunostaining for phospho-Histone H3 (Ser10) was performed. Phosphorylation of Histone H3 (pHH3) at Ser10 can be observed during mitosis in all eukaryotes and is therefore a reliable marker for cells under-going division^83,84^. All embryos were counter-stained with DAPI (1:1000). Samples were mounted with the anterior endoderm facing the coverslip, and only the first 50 z-slices (100 *µ*m) were analysed. Nuclei positive for TUNEL or phospho-Histone H3 (Ser10) were manually counted using the “Cell Counter” plugin in ImageJ or the ‘spot’ function in Imaris. To allow easier discrimination between endoderm and the underlying mesoderm/epiblast cells, images were re-sliced in the proximal-distal direction, and only cells in the outermost cell layer were considered for quantification. Coordinates relative to the centre of invagination were normalised and plotted as described above for live imaging data sets.

#### Analysis of crowdedness in live imaging datasets

To visualise the level of crowdedness surrounding the involuting foregut over the involution stages we first utilised an existing tracking dataset^37^ which was generated from a ubiquitously-labelled nuclear marker^21^. This dataset was loaded into Imaris and the nearest 5 neighbours were computed and visualised along a colour scale corresponding to the degree of crowdedness. To determine the crowdedness of the endoderm layer over the course of foregut pocket formation, live imaging data sets with nuclear labelling of all endoderm cells (*Foxa2*-eGFP line) were used. Nuclei were automatically labelled using the “spot detection” function in Imaris, followed by manual correction of the results. The distances to the nearest 5 neighbours, computed by Imaris, were exported as a .csv file and used for further analysis of cell densities in the endoderm.

#### Analysis of cCasp3 staining

Embryos were stained for cleaved caspase-3 (cCasp3) as described earlier and counter-stained with DAPI (1:1000). After immunostaining, samples were mounted with the anterior endoderm facing the coverslip, and the first 50 z-slices (100 *µ*m) were analysed. Cells positive for cCasp3 were manually counted using the “Cell Counter” plugin in ImageJ or the ‘spot’ function in Imaris. To allow easier discrimination between endoderm and the underlying mesoderm/epiblast cells, images were re-sliced in the proximal-distal direction, and only cells in the outermost cell layer were considered for quantification. To discriminate between emVE and DE cells within the endoderm layer, embryos expressing *Afp*-kGFP were used, and emVE cells were identified by GFP expression.

#### Statistical analyses

Embryos were randomly allocated to treatment groups in culture experiments, and the investigator was not blinded to group allocation. Sample size was determined based on previous experimental experience and ethical considerations, to minimise the number of animals used while ensuring statistical validity. All statistical analyses in this study, with the exception of RNA sequencing analysis, were performed using GraphPad Prism. Data were tested for normal distribution using a Kolmogorov-Smirnov test. Data that did not show a Gaussian distribution were analysed using a Mann-Whitney U test (two groups) or a Kruskal-Wallis test (multiple groups). Normally distributed data were analysed with an unpaired two-tailed Student’s t-test (two groups) or an ANOVA test (multiple groups). A Welch’s correction was applied if the variances between the groups were significantly different. All quantitative data are shown as mean *±* SD. Detailed information about the statistical test used can be found in the corresponding figure legend.

### Morphometric analysis

#### Mesh generation

To produce a closed triangular mesh covering the embryo surface we used a shrink-wrap algorithm. We started from the raw light-sheet volumetric images: each image volume was converted into a point cloud by thresholding at the mean intensity plus two standard deviations and extracting the perimeter of the resulting binary image with bw-perim in MATLAB^85^. We then filled the embryonic cup with synthetic points by computing the convex hull of the point cloud and generating points at a fixed distance (20*µm*) inside its surface; these interior points prevent the shrink-wrap from collapsing into the concave cup. To increase resolution, we added further triangulation to the convex hull and applied the shrink-wrap algorithm to tighten the mesh onto the embryonic point cloud. Finally, we recomputed a mesh from the shrink-wrapped vertices using the pc2facemesh function in MATLAB, and applied a series of mesh-healing operations with the Python package PyMesh-Lab^86^. Code used to produce these results is publicly available here: github.com/gserranonajera/MouseFlowshape.

#### Spherical harmonic computation

Shape descriptors were computed from each closed triangular mesh after masking out the posterior and proximal regions to isolate the folding of the foregut. This step also avoids introducing sharp edges along the proximal rim of the embryonic cup or from areas that fell out of view in some experiments, ensuring that all embryos present comparable features. Spherical harmonic decomposition was performed using FlowShape^22^ up to degree *ℓ =* 15 (16 modes), giving 256 coefficients per shape.

#### Shape trajectory analysis

For each embryo and timepoint, we computed the spherical harmonic power spectrum from the masked coefficients and summed the power over degrees *ℓ* ≥ 2 to obtain a single scalar measure of shape complexity; the *ℓ =* 0 and *ℓ =* 1 terms were excluded so that the measure reflects shape rather than overall scale or position. This measure was smoothed over time with a running median (window = 5 frames, 25 min), and outliers were removed using an interquartile-range criterion. The time of maximum involution was defined as the peak of the smoothed complexity curve and used as a temporal landmark to align embryos. To place all embryos in a common frame of reference, each embryo’s coefficients were rotated into a shared orientation directly in the spherical harmonic domain using FlowShape and manual adjustments if necessary. Rotations were anchored at the first and peak timepoints and interpolated between them by spherical linear interpolation (SLERP) implementation included in scipy library^87^, holding the peak orientation for later timepoints. Finally, principal component analysis (PCA) was fit on the control embryos only, and mutant embryos (*Bcl2*-OE, *Bmp2*-VE-KO) were projected into the resulting space. The first two principal components were used to visualise the morphogenetic trajectory of foregut involution over developmental time.

## Supplementary Information

### Supplementary Data Figures

**Supplementary Figure 1:**
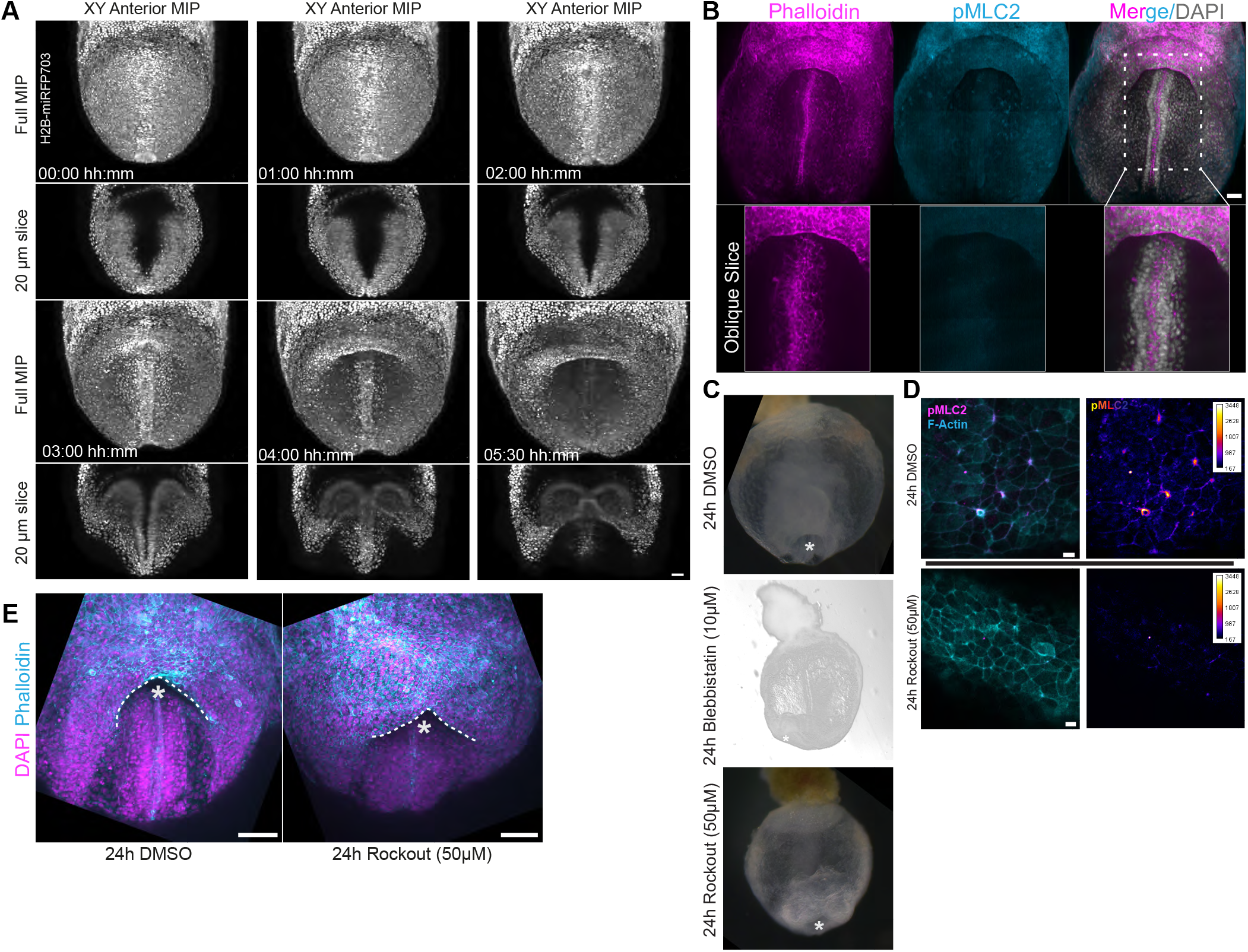
Progression of foregut development and actomyosin inhibition. **A** Still images from light-sheet imaging of an H2B-miRFP703 embryo from around the LB stage. MIP projections of the anterior region and 20 *µ*m headfold sections. Scale bar 40 *µ*m. **B** pMLC and Phalloidin staining of an LHF stage embryo, showing actin condensation over the notochord but no pMLC signal. Zoomed-in view of 85 *µ*m thick oblique slices. Scale bar 50 *µ*m. **C** Widefield images of the anterior view of E8.5 embryos after 24 h of *ex vivo* culture in DMSO, 10 *µ*M Blebbistatin, or 50 *µ*M Rockout. The position of the foregut is indicated by an asterisk. **D** Wholemount immunostaining for F-actin (cyan) and pMLC2 (magenta) of E8.5 embryos after 24 h culture with DMSO or 50 *µ*M Rockout. Sum projection of 5 z-slices (z-slice thickness: 2 *µ*m). Scale bar 10 *µ*m. **E** Representative DMSO-treated control and 50 *µ*M Rockout-treated embryos. Involution still proceeds under inhibitor treatment but forms a shallower, more pointed pocket (* marks the anterior tip). Scale bar 100 *µ*m.

**Supplementary Figure 2:**
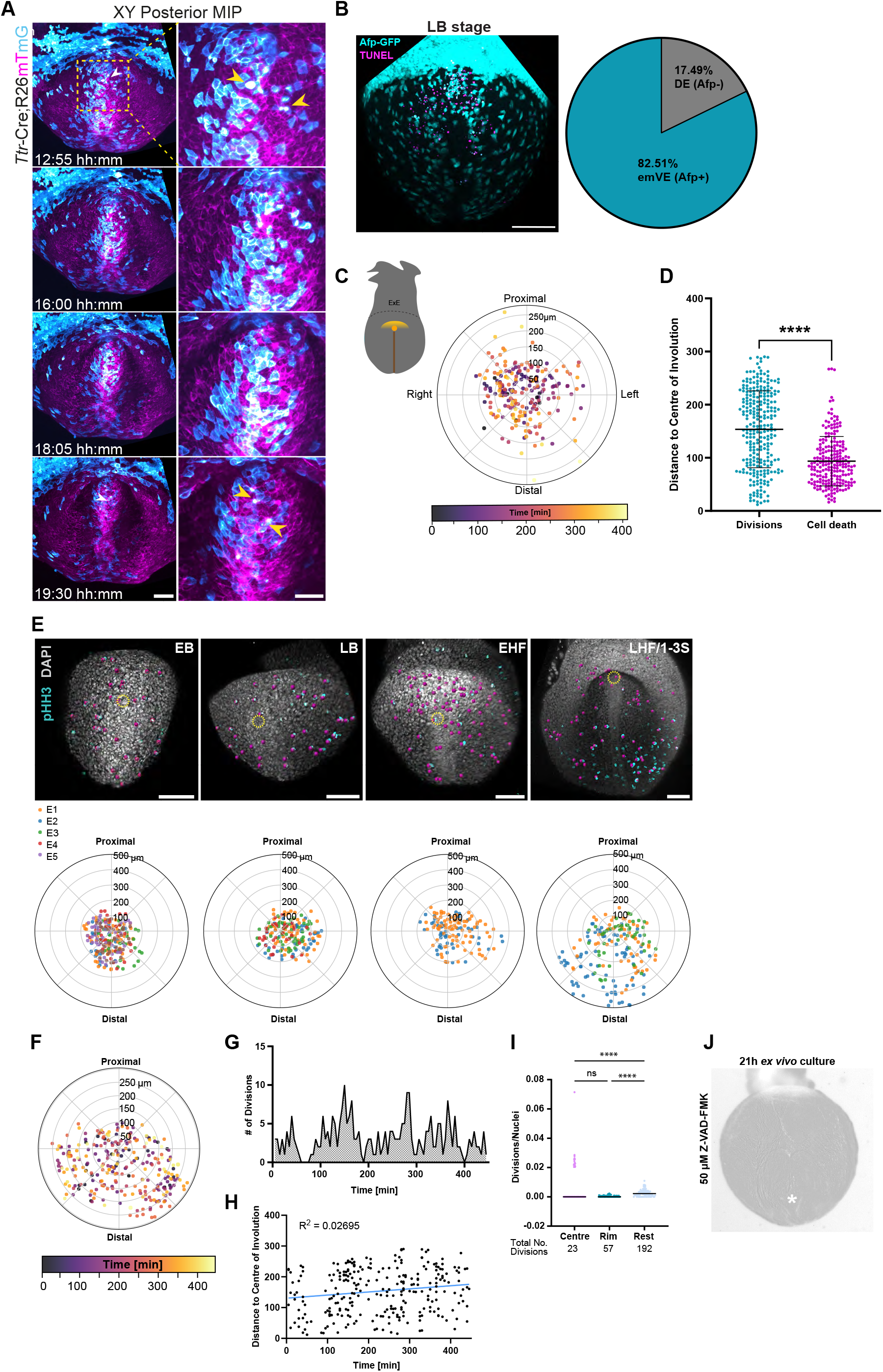
Localisation of cell death in the posterior and rates of proliferation in the anterior. **A** Selected posterior views and zoomed-in insets of the boxed regions (yellow dashed box; right panels) from a live imaging data set of a representative *Ttr*-Cre^*TG/+*^; Rosa26R^*mTmG*^ embryo (the same embryo shown in Figure 3A). Arrows point to visceral endoderm cells showing signs of cell death, indicated by blebbing and extrusion from the epithelium. Maximum intensity projections. Scale bars 100 *µ*m (left panels), 50 *µ*m (right panels). **B** Anterior view of a late bud stage *Afp*-kGFP (cyan) embryo subject to the TUNEL assay (magenta) to detect cell death. The pie chart shows quantification of death events in GFP-positive or negative cells in the endoderm layer. Scale bar 100 *µ*m. N = 4 embryos (LB stage), n = 749 TUNEL-positive events. **C** Quantification of the distance of cell death events relative to the centre of invagination (orange dot in schematic, centre of rose plot), colour-coded by time. N = 2 embryos, n = 218 death events. Colour-coded by time in minutes. **D** Analysis of the distance of cell division and death events relative to the centre of invagination quantified in a live imaging dataset. N = 1 embryo (divisions), 2 embryos (cell death); n = 272 (division events), 218 (cell death events). Mann-Whitney test. P-value: **** ≤ 0.0001. Mean and SD. Cell death data is colour-coded by replicate. **E** Representative images of E7.5 mouse embryos at different stages, immunostained for phospho-Histone H3 (Ser10) (pHH3, cyan) and DAPI (grey). pHH3-positive cells in the outer endoderm layer are labelled with magenta dots, and the centre of the future foregut pocket is indicated with a yellow circle. Scale bars 100 *µ*m. Rose plots corresponding to each stage are shown underneath. The orientation of the rose plots corresponds to the orientation of the representative samples above, with the centre of the rose plot as the centre of foregut involution. Dots are colour-coded by replicate. N = 6 (EB), 4 (LB), 2 (EHF), 3 (LHF) embryos; n = 224 (EB), 181 (LB), 131 (EHF), 187 (LHF) divisions. **F** Analysis of cell divisions in the anterior endoderm from a time-lapse of a *Foxa2*-eGFP-expressing embryo. The centre of the rose plot corresponds to the centre of foregut invagination. Division events are colour-coded by time. N = 1 embryo, n = 272 division events. **G–I** Analysis of the number of divisions over time (**G**), the distance of divisions relative to the centre of involution (**H**), and the number of divisions in the centre, rim, and rest region relative to the total number of nuclei in each region at any given time point (**I**), of a representative time-lapse imaging of a *Foxa2*-eGFP-expressing embryo also used for **F. J** Anterior view of a representative E7.5 mouse embryo cultured for 21 hours with pan-caspase inhibitor Z-VAD-FMK (50 *µ*M). The foregut pocket is indicated by an asterisk.

**Supplementary Figure 3:**
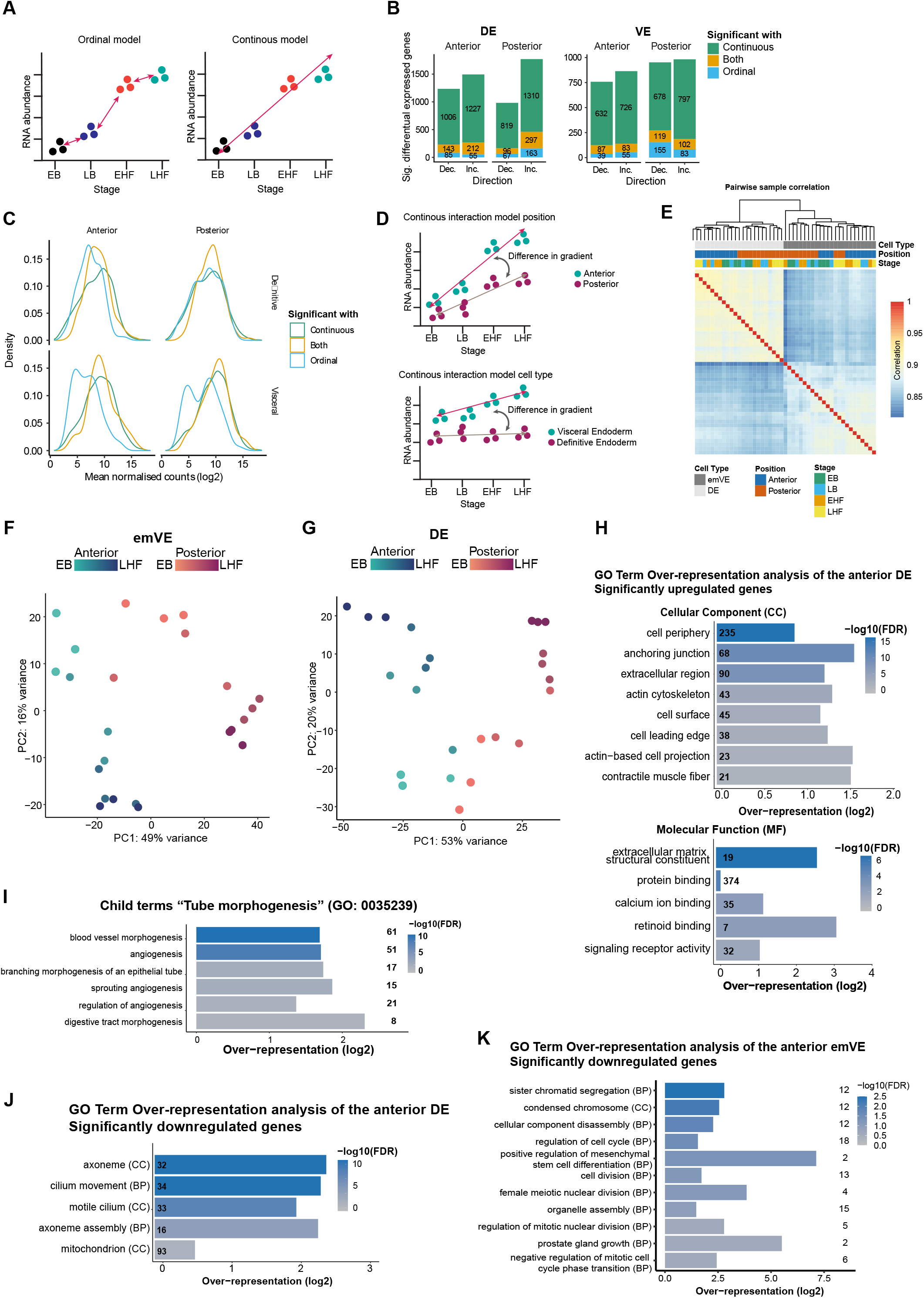
Models for statistical testing of differential gene expression and gene comparison. **A** Schematic illustration of the ordinal and continuous main effects models for statistical testing. **B** Comparison of significantly differentially expressed genes detected using the continuous or ordinal main effects models. **C** Comparison of the density and read counts of differentially expressed genes in the continuous and ordinal models. **D** Schematic illustration of modifications of the continuous model to account for positional and cell type information by introducing interaction terms. **E** Exploration of sample separation between DE and emVE samples using pairwise comparisons. DE and emVE cluster by position and stage. **F–G** PCA plots of the emVE and DE over the different developmental stages of E7.5. **H** Over-represented GO terms among significantly upregulated genes in the anterior DE vs. anterior emVE and posterior DE, by Cellular Component (CC) and Molecular Function (MF). **I** Over-represented child terms of GO:0035239, “tube morphogenesis”, among the significantly overexpressed genes in the anterior DE. Adjusted *P <* 0.01. **J** Over-represented GO terms among the significantly downregulated genes in the anterior DE relative to the anterior emVE and posterior DE. Adjusted *P <* 0.01. Numbers in the bars indicate the number of differentially expressed genes belonging to each GO term. **K** Over-represented GO terms in the gene set of significantly downregulated genes in the anterior emVE relative to the anterior DE and posterior emVE. No over-represented GO terms were detected in the upregulated gene set. Numbers to the right indicate the number of differentially expressed genes belonging to each GO term. Adjusted *P <* 0.01.

**Supplementary Figure 4:**
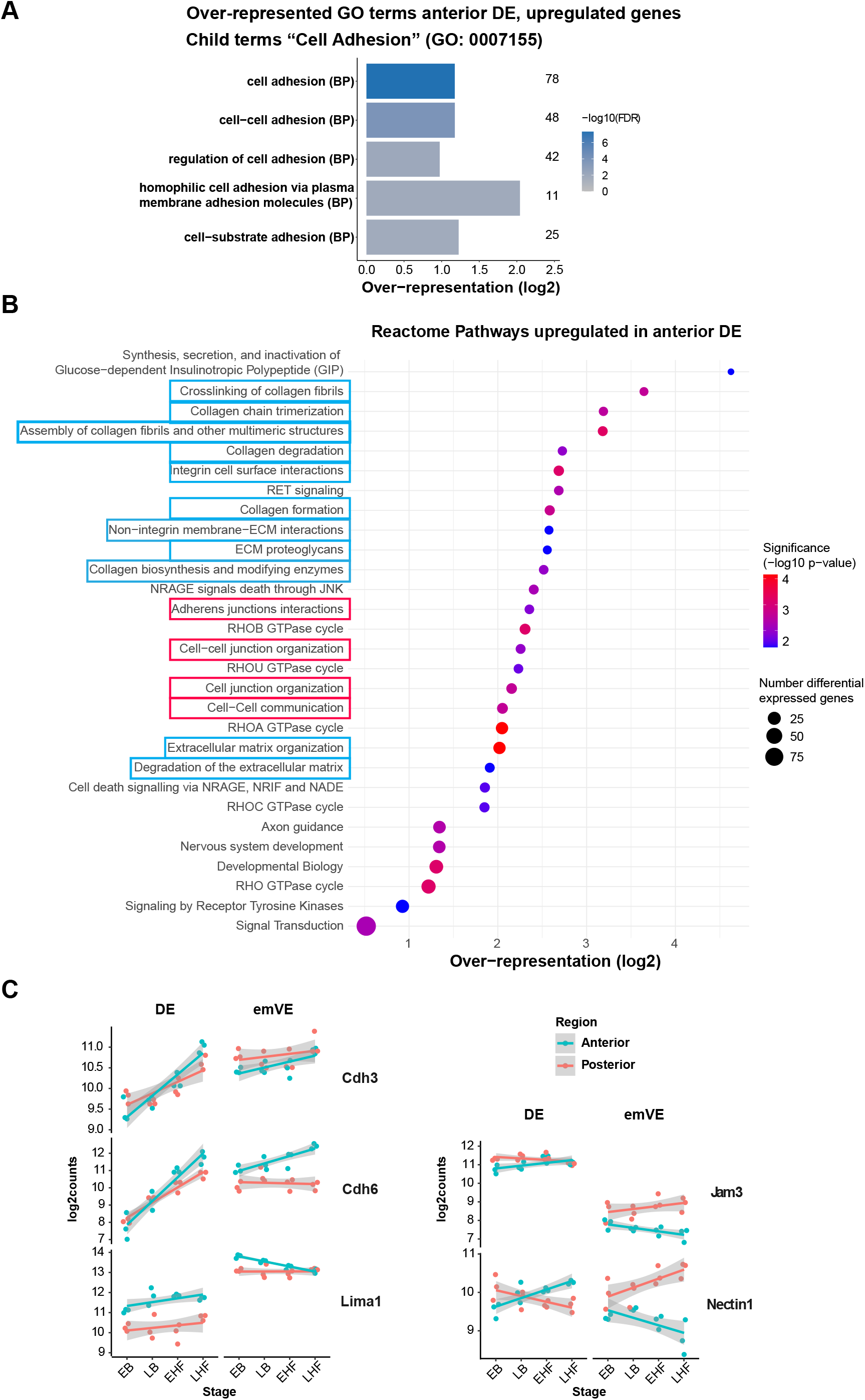
Exploration of differential expression of genes related to cell adhesion. **A** GO-term exploration of the GO term “cell adhesion” (GO:0007155) and its child terms in the upregulated gene set of anterior DE cells. Numbers to the right indicate the number of differentially expressed genes belonging to each GO term. **B** Reactome pathway analysis of the significantly upregulated genes in anterior DE cells. Cell-cell adhesion terms are highlighted in red, and cell-matrix adhesion terms in blue. **C** Selected genes associated with cell adhesion showing significant differential expression between DE and emVE cells. Dots represent individual replicates; lines show the fitted linear regression with confidence interval.

**Supplementary Figure 5:**
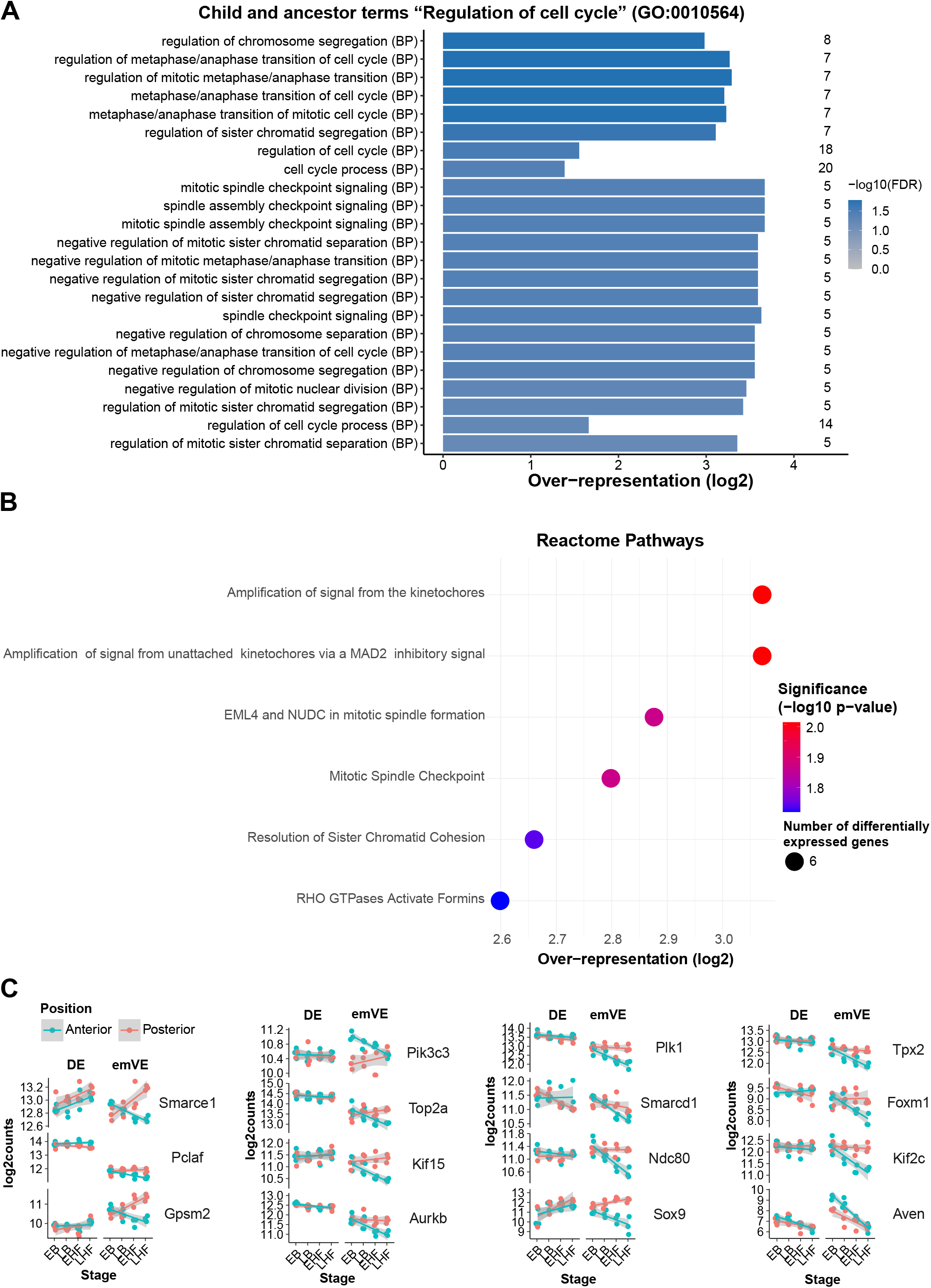
Down-regulation of cell cycle genes in the anterior emVE. **A** Child and ancestor terms of the GO term “sister chromatid segregation” (GO:0000819) over-represented among the significantly down-regulated genes in the anterior emVE compared to the posterior emVE and anterior DE. Adjusted *P <* 0.01. Numbers to the right indicate the number of differentially expressed genes belonging to each GO term. **B** Over-represented Reactome pathways among the genes significantly downregulated in the anterior emVE. **C** Selected differentially expressed genes associated with the GO term “cell cycle process” in the downregulated gene set of anterior emVE cells. Dots represent individual replicates; lines show the fitted linear regression, with the confidence interval shown in grey.

**Supplementary Figure 6:**
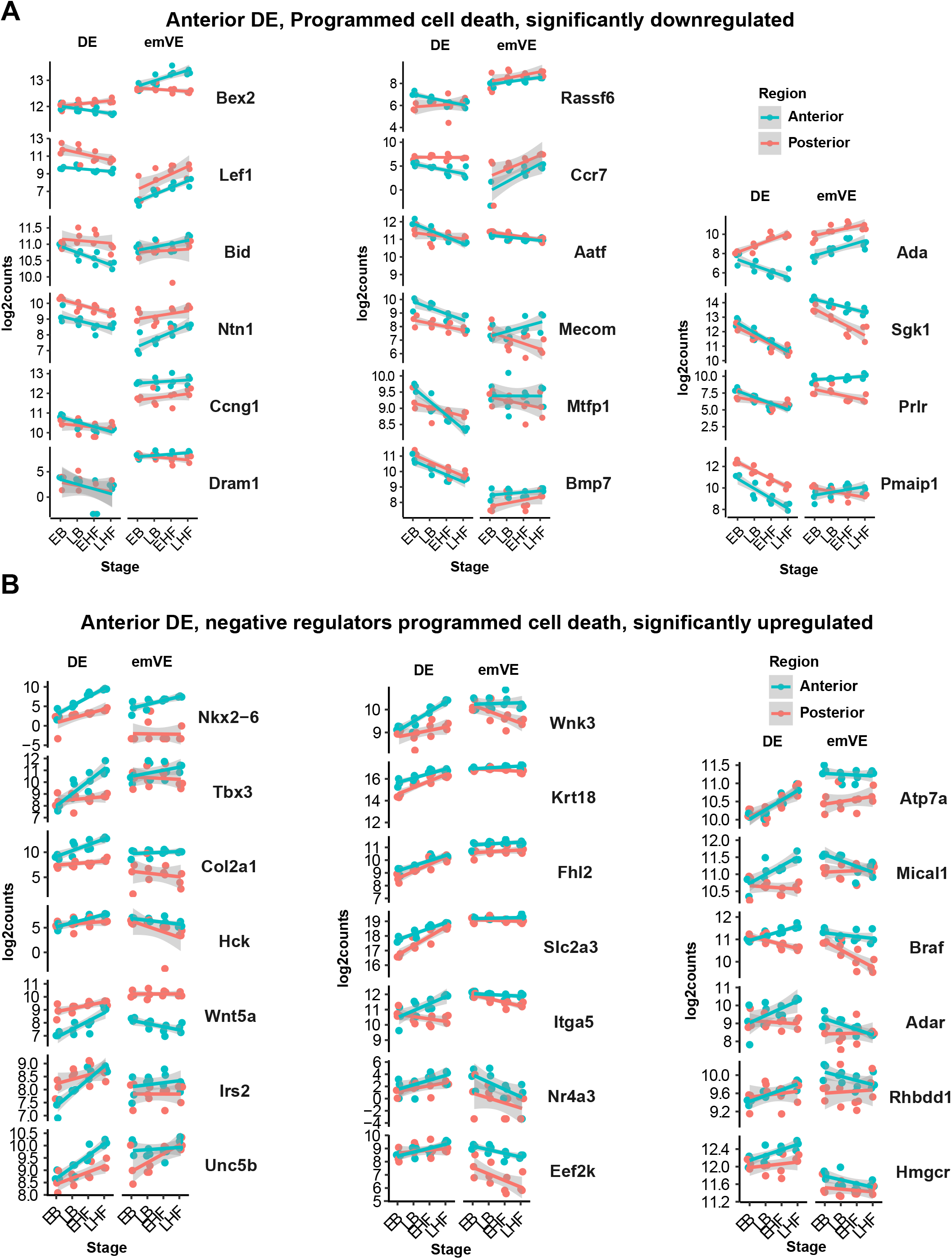
Differential expression of anti- and pro-apoptotic genes in the anterior DE. **A** RNA expression of genes significantly downregulated relative to anterior emVE in the anterior DE, associated with “programmed cell death” (GO:0012501). **B** RNA expression of genes significantly upregulated relative to anterior emVE in the anterior DE, associated with “negative regulation of programmed cell death” (GO:0043069). Dots represent individual replicates; lines show the fitted linear regression with confidence interval.

**Supplementary Figure 7:**
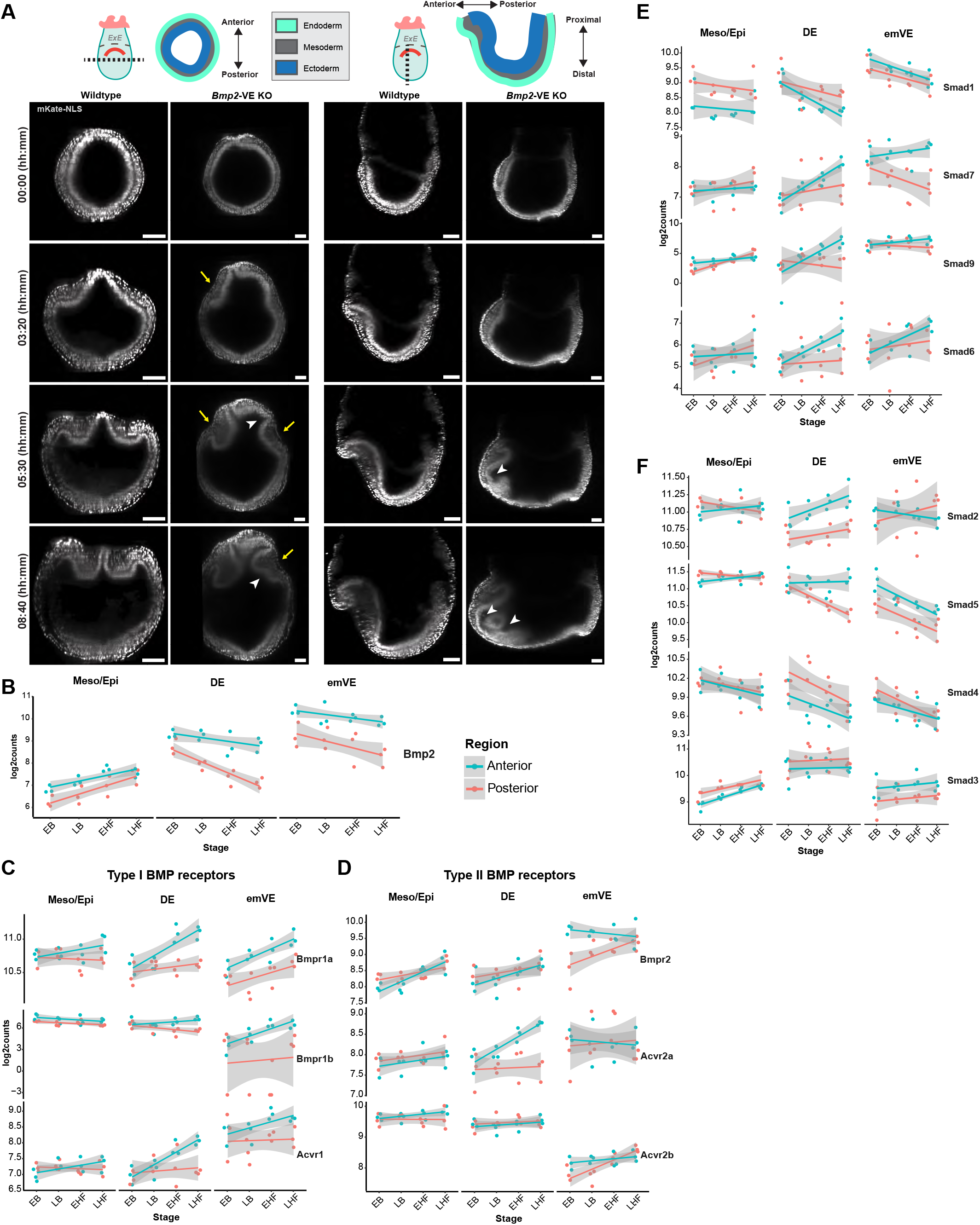
The role of BMP during foregut involution. A Comparison of the development of a WT and Bmp2-VE KO embryo expressing mKate2-nls. Shown are selected orthogonal views along the y -z (left panels) and x-y (right panels) axis from a time-lapse imaging. The positions of the image slices as well as the orientation of the embryo are indicated in the pictograms at the top. Yellow arrows point to the forming lateral invaginations in the mutant, white arrowheads indicate ectopic folding of the headfolds. Scale bars 100 µm. B Bmp2 expression in the anterior and posterior emVE, DE, and mesoderm/epiblast (Mes/Epi) cell populations of the E7.5 mouse embryo. C–D Gene expression levels of type I and type II BMP receptors. E–F Gene expression levels of different receptor-regulated and inhibitory Smad proteins as downstream effectors of the canonical BMP-signalling pathway. Dots represent individual replicates; lines show the fitted linear regression with confidence interval.

**Supplementary Figure 8:**
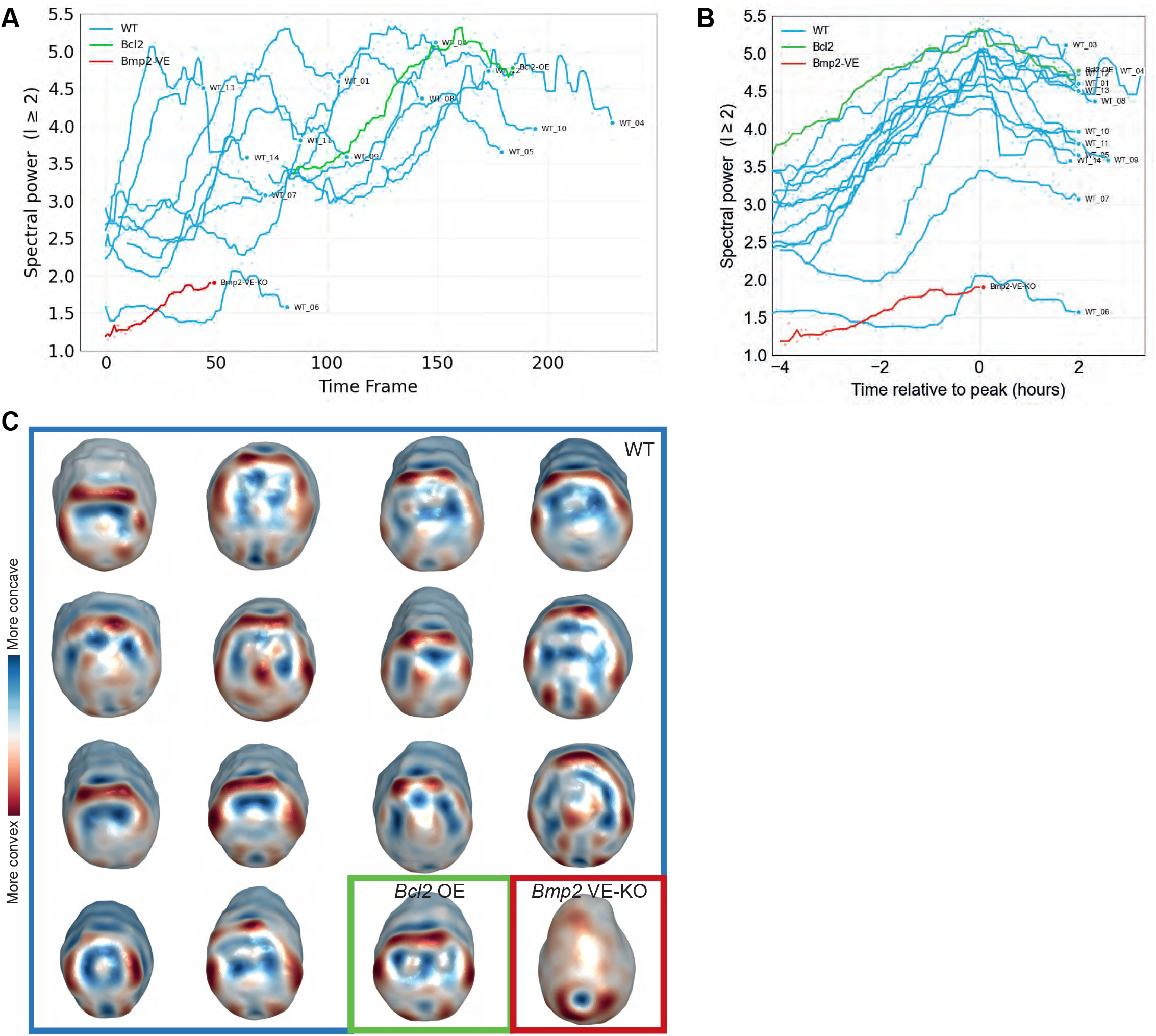
Morphometric analysis of wild-type and mutant embryos. **A** Shape complexity over time. Complexity is measured by the spectral power: the total energy in the spherical harmonic coefficients at each time point. Higher values indicate more shape complexity. Wild-type embryos show a stereotyped peak coinciding with maximal curvature during involution while the *Bmp2*-VE KO mutant has no maximal peak. **B** Temporal alignment of morphogenetic trajectories using the spectral powers from (A). **C** Meshes representing the point of maximum complexity across fourteen different WT embryos, as well as *Bcl2*-VE over-expression and *Bmp2*-VE KO mutants.

**Table S1:** Overview of embryo collection and RNA-sequencing sample sizes. Embryos were collected at four developmental sub-stages (EB, LB, EHF, LHF; 112 embryos total), dissected into anterior and posterior halves, and dissociated and sorted by FACS into emVE (*Afp*-kGFP^+^/*Foxa2*-eGFP^+^), DE (kGFP^−^/eGFP^+^), and Mes/Epi (kGFP^−^/eGFP^−^) populations. Cells from multiple embryos of the same stage, position, and population were pooled and split into biological triplicates prior to RNA extraction and sequencing; pooled and mean per-replicate cell counts are shown.

| Stage | N embryos | Position | Population | Cells pooled | Mean cells/replicate (n=3) |
| --- | --- | --- | --- | --- | --- |
| EB | 34 | Anterior | emVE | 3,045 | 1,015 |
|  |  |  | DE | 6,988 | 2,329 |
|  |  |  | Mes/Epi | 42,357 | 14,119 |
|  |  | Posterior | emVE | 1,059 | 353 |
|  |  |  | DE | 5,673 | 1,891 |
|  |  |  | Mes/Epi | 44,338 | 14,779 |
| LB | 30 | Anterior | emVE | 2,960 | 987 |
|  |  |  | DE | 10,343 | 3,448 |
|  |  |  | Mes/Epi | 38,920 | 12,973 |
|  |  | Posterior | emVE | 1,382 | 461 |
|  |  |  | DE | 5,638 | 1,879 |
|  |  |  | Mes/Epi | 41,333 | 13,778 |
| EHF | 27 | Anterior | emVE | 2,286 | 762 |
|  |  |  | DE | 11,154 | 3,718 |
|  |  |  | Mes/Epi | 46,601 | 15,534 |
|  |  | Posterior | emVE | 747 | 249 |
|  |  |  | DE | 3,065 | 1,022 |
|  |  |  | Mes/Epi | 45,000 | 15,000 |
| LHF | 21 | Anterior | emVE | 3,792 | 1,264 |
|  |  |  | DE | 18,096 | 6,032 |
|  |  |  | Mes/Epi | 40,000 | 13,333 |
|  |  | Posterior | emVE | 2,214 | 738 |
|  |  |  | DE | 6,864 | 2,288 |
|  |  |  | Mes/Epi | 39,812 | 13,271 |

### Movies

**Movie S1** Anterior maximum intensity projection time-lapse of an H2B-miRFP703 expressing mouse embryo starting at the LB stage imaged using light-sheet microscopy. Scale bar 50 *µ*m, time interval 5 minutes.

**Movie S2** Live-imaging of a LifeAct-RFP embryo from LB to LHF stage using light-sheet microscopy, showing actin cytoskeleton changes over the lip of the foregut pocket and condensation over the notochord. Maximum intensity projection of the anterior side. Scale bar 50 *µ*m, time interval 5 minutes.

**Movie S3** Three-dimensional mesh computed from live-imaging data of a LB to EHF mouse embryo using light-sheet microscopy. Color indicates degree of curvature, red being more convex, blue more concave.

**Movie S4** Cell density of an H2B-eGFP embryo imaged with light-sheet microscopy; computed whole-embryo tracks and lineages from^37^. The anterior-most proximal region and the midline become progressively more crowded over time (average distance to 5 nearest neighbours; blue = crowded, red = sparse). Scale bar 50 *µ*m, time interval 5 minutes.

**Movie S5** Anterior MIPs from live imaging using light-sheet microscopy of a *Ttr*-Cre;ROSA26^mTmG^ embryo. Scale bar 50 *µ*m, time interval 5 minutes.

**Movie S6** Live imaging using spinning disk microscopy of an E7.5 embryo expressing *Afp*-kGFP (green) and LifeAct-RFP (magenta and grey, right panel; 3×3×1 median filter), showing emVE cell extrusion and formation of a contractile actomyosin ring. Scale bar 3 *µ*m, time interval 3 minutes.

**Movie S7** *Bcl2*-VE Over-expression embryo imaged using light-sheet microscopy. Embryo is ubiquitous for myr-eGFP, LUT mpl-inferno. Scale bar 50 *µ*m, time interval 5 minutes.

**Movie S8** Light-sheet imaging of a *Bmp2*-VE KO embryo expressing mKate2-nls (cyan; anterior left, posterior right, YZ Projection). Wavelet fusion, 50 *µ*m size, 1 pixel median filters. Scale bar 50 *µ*m, time interval 5 minutes.

